# Predicting TCR-epitope Binding Specificity Using Deep Metric Learning and Multimodal Learning

**DOI:** 10.1101/2021.03.19.436191

**Authors:** Alan M. Luu, Jacob R. Leistico, Tim Miller, Somang Kim, Jun S. Song

## Abstract

Understanding the recognition of specific epitopes by cytotoxic T cells is a central problem in immunology. Although predicting binding between peptides and the class I Major Histocompatibility Complex (MHC) has had success, predicting interactions between T cell receptors (TCRs) and MHC class I-peptide complexes (pMHC) remains elusive. This paper utilizes a convolutional neural network model employing deep metric learning and multimodal learning to perform two critical tasks in TCR-epitope binding prediction: identifying the TCRs that bind a given epitope from a TCR repertoire, and identifying the binding epitope of a given TCR from a list of candidate epitopes. Our model can perform both tasks simultaneously and reveals that inconsistent preprocessing of CDR3B sequences can confound binding prediction. Applying a neural network interpretation method identifies key amino acid sequence patterns and positions within the TCR important for binding specificity. Contrary to the common assumption, known crystal structures of TCR-pMHC complexes show that the predicted salient amino acid positions are not necessarily the closest to the epitopes, implying that physical proximity may not be a good proxy for importance in determining TCR-epitope specificity. Our work thus provides insight into the learned predictive features of TCR-epitope binding specificity and advances associated classification tasks.

## INTRODUCTION

The human adaptive immune response requires a two-step binding process to mediate the recognition and destruction of diseased cells by Cytotoxic T lymphocytes (CTLs). First, the class I MHC molecules found in all nucleated cells bind select peptides resulting from cleaving cytosolic proteins, some of which may derive from pathogens infecting the host cells, and present them on the cell surface as a peptide-MHC (pMHC) complex. Second, the T cell receptors (TCRs) of CTLs must scan the presented pMHCs and bind only those pMHCs containing specific peptides recognized by the TCRs. Most human CTLs contain TCRs consisting of α and β glycoprotein chains, and only a small subset of CTLs contain TCRs composed of γ and δ chains. During the binding interaction between the TCR and pMHC, the Complementarity-Determining Region 1 (CDR1) and Complementarity-Determining Region 2 (CDR2) loops of the TCR chains typically make significant contact with the MHC, while the Complementarity-Determining Region 3 (CDR3) loop directly interacts with the peptide itself and thus determines the specificity of the TCR and pMHC interaction (1). Peptides recognized by the TCR in this context are known as epitopes. Currently, a number of immune assays are used to measure T cell response to pathogen peptides (2–6). Researchers have attempted to predict both peptide-MHC and TCR-epitope binding events using the amino acid sequence information contained in experimental TCR-pMHC binding data emerging from recent TCR sequencing technology (7,8). Notable progress has been made in predicting binding between peptides and MHC (9–12), but the peptide-MHC binding is only a necessary, not sufficient, condition for TCR-epitope binding (13), the prediction of which has had only limited success.

A variety of approaches to TCR-epitope binding prediction have been explored in the literature. For instance, Glanville et al. (14) and Dash et al. (15) have utilized amino acid motif signatures present in the CDR3 region of the TCR α and β chains to cluster TCRs by epitope specificity. Methods modeling the physical structure and interaction between TCR and pMHC have also been proposed (16–19). Additionally, with the emergence of large databases of aggregated experimental TCR-epitope binding data, researchers have begun to explore machine learning methods, including Gaussian Processes (20), convolutional neural networks (21,22), and recurrent neural networks (23). These methods differ in terms of datasets used for training and testing, as well as the nature of the classification tasks performed, making direct comparison of model performance difficult.

Ideally, prediction of TCR-epitope binding would allow classifying any pair of TCR and pMHC into a binding or non-binding class given 3 inputs: TCR variable region sequence, peptide sequence, and the HLA allele of the MHC molecule. However, limitations of data availability and theoretical understanding currently restrict this task to only special considerations relying on certain assumptions. First, previous investigations have analyzed TCR-pMHC crystal structures and shown that the CDR3 loop of TCR α chain (CDR3A) rarely makes contact with the epitope, whereas the CDR3 loop of TCR β chain (CDR3B) always makes contact with the epitope, suggesting that the CDR3B sequence may play a dominant role in determining the TCR-epitope binding specificity (14). Consequently, most large databases reporting TCR-pMHC interactions include only the CDR3B sequences, and prediction methods utilizing large amounts of data primarily use only this limited information, as is the case for this manuscript. Second, most databases only report confirmed instances of TCR-pMHC binding, known as positive binding data, and lack confirmed non-binding events between a TCR and pMHC, known as negative binding data, which are also needed for robust TCR-epitope binding prediction. Previous studies (20–23) have generated artificial negative binding data through two approaches that assume a TCR and an epitope not explicitly confirmed to bind are non-binding: one approach is to utilize only positive binding data and randomize TCR pairing with reported epitopes; another approach, which is employed in this manuscript, utilizes an ambient set of TCRs obtained via high-throughput sequencing of the human TCR repertoire and not present in the positive set, assuming that every TCR in this ambient set does not bind an epitope in the positive set. Third, currently available TCR-pMHC binding data lack diversity in the HLA allele and epitope sequence, with the vast majority of reported interactions associated with a small number of HLA alleles and peptides. Some studies have addressed this issue by limiting their analysis to TCR-pMHC binding interactions involving the most common HLA allele (21), HLA A*02:01, as also pursued in this work. The lack of epitope diversity implies that TCRs and epitopes usually display a many-to-one relationship, with many distinct TCRs binding each curated epitope. Cross-reactivity of TCRs (24), the phenomenon of a single TCR recognizing multiple epitopes, is also reported in public databases, but is relatively rare. Consequently, current models are usually capable of predicting binding between an epitope the model has seen during training and a new TCR previously unseen during training, but tend to fail at predicting binding for a new epitope unseen during training. Because of this limitation, most studies currently attempt to predict binding only between a small predetermined set of epitopes and arbitrary TCRs, rather than between arbitrary epitopes and TCRs.

This manuscript formalizes the above limitations, which were only implicitly addressed in most previous studies, by categorizing the TCR-epitope binding prediction problem into 3 distinct tasks. Task 1 involves identifying the specific TCRs that can bind a given epitope sequence from a repertoire of TCRs. This task can be considered as a binary classification of TCRs into binding and non-binding classes. The appropriate performance metric for this task is the Area Under the ROC Curve (AUC). Task 2 involves identifying the binding epitope of a TCR from a predetermined list of candidate epitopes. This task can be considered as a multi-class classification problem of classifying TCRs into the candidate epitope-binding groups. The appropriate performance metric in this setting is the classification accuracy. Finally, Task 3 involves predicting whether any arbitrary TCR and epitope pair will interact; this task is the most complex and is currently infeasible because of the lack of relevant training data, as only few epitopes currently have reported interactions with a large number of unique TCRs in the available databases of TCR-epitope binding. We here propose a novel convolutional neural network (CNN) model, inspired by ideas from deep metric learning (25–27) and multimodal learning (28), directly utilizing the CDR3B sequences of TCRs as well as the epitope sequences. Even though the model is, in principle, capable of simultaneously performing all three prediction tasks given sufficient data, this manuscript shall focus on implementing and testing the model only for Task 1 and Task 2, as there currently do not exist sufficient data required for Task 3.

Our model requires only a single training procedure for all three tasks, but evaluation on each of the three tasks can be performed by modifying the composition of the test set and the procedure for evaluating model output. The architecture of our CNN model revolves around the observation that the TCR-epitope binding prediction problem may be considered a multimodal machine learning problem. Modality in machine learning is analogous to *sensory modality*, defined as our “primary channel of communication and sensation, such as vision or touch” (28). In the setting of machine learning, modalities correspond to classes of data types, such as image data and text data. Multimodal machine learning therefore involves modeling the interactions between multiple data types. We thus consider TCR-epitope binding to be a multimodal machine learning problem, with the CDR3B sequences and epitope sequences considered as two separate modalities. A central design choice in multimodal machine learning is the choice of representation of the multimodal input. In this aspect, our work differs from previous studies utilizing neural networks (21,23) based on a joint representation scheme in which CDR3B and epitope sequences are combined into a single representation as model input. By contrast, our model uses a coordinated representation scheme in which the CDR3B and epitope sequence inputs, initially encoded in a numeric matrix form (Figure 1A; Methods: CDR3B and epitope sequence representation), are each mapped to their own representation in a latent vector space by a CDR3B embedding network and epitope mapping network, respectively (Supplementary Figure S1,S2; Supplementary Methods: Model architecture and training). The latent space representations of the CDR3B and epitope sequences are then coordinated through a similarity measure using a composite neural network model comprising two subunits: the Triplet Network (25,26) and Modal Alignment Network (Figure 1B; Supplementary Methods: Model architecture and training). The Triplet Network itself is a composite structure that trains the CDR3B embedding network to map CDR3B sequences to points in a latent space, clustered into point clouds according to their binding epitope, by minimizing Triplet Loss (Figure 1C, Supplementary Figure S3; Supplementary Methods: Model architecture and training); intuitively, training the Triplet Network to minimize Triplet Loss forces the network to pull together the points corresponding to CDR3B sequences binding the same epitope, while pushing apart the points corresponding to CDR3B sequences binding different epitopes. Another neural network, known as the Modal Alignment Network (Figure 1D, Supplementary Figure S4; Supplementary Methods: Model architecture and training), then maps epitope sequences to Gaussian distributions localized to these point clouds in the aforementioned latent space. We demonstrate that this model performs similarly to, and in some cases outperforms, current state-of-the-art methods, while providing a flexible computational framework for simultaneously handling multifarious classification tasks.

**Figures 1:**
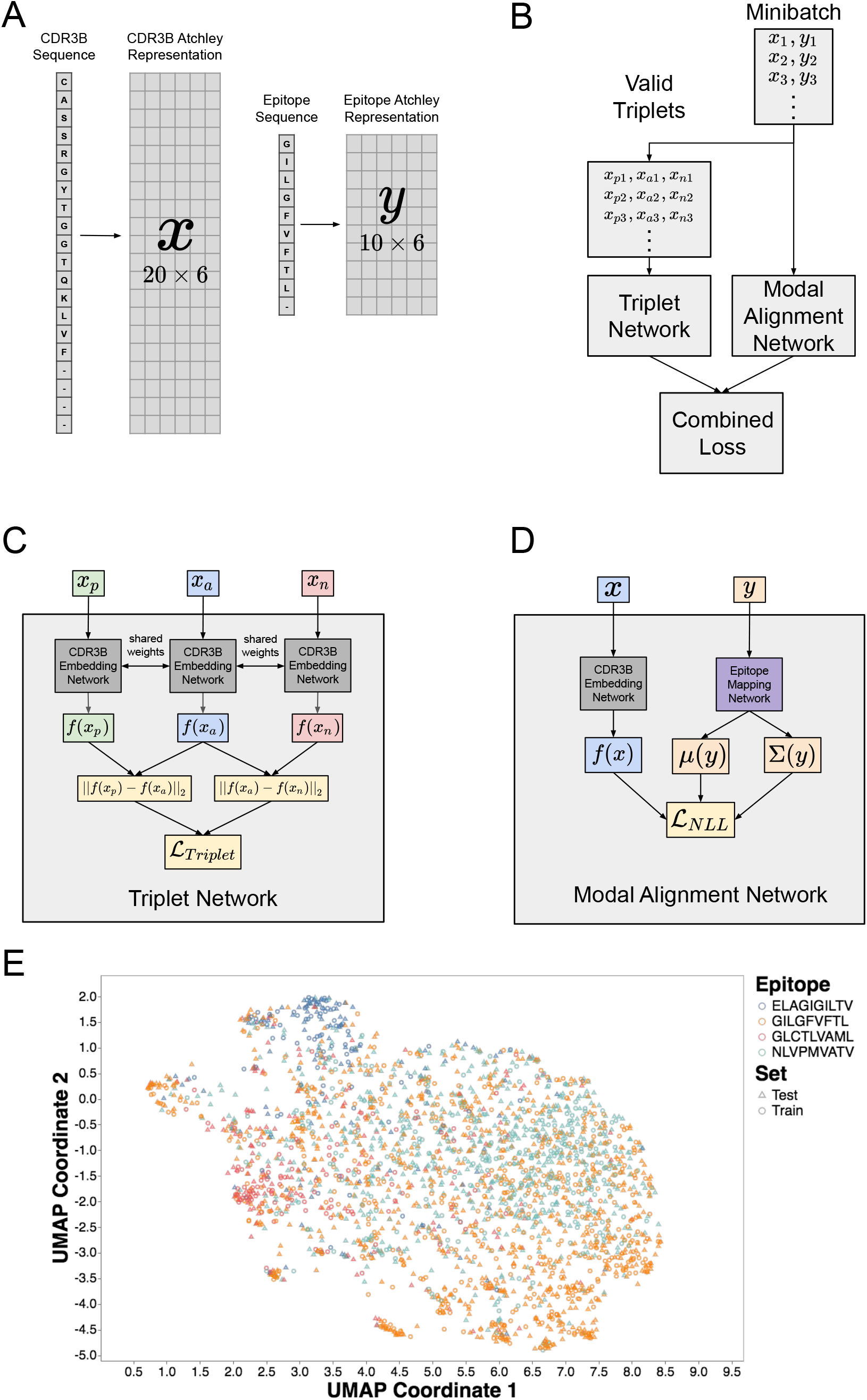
Amino acid sequence representation and architecture of hybrid neural network model. **(A)** Visualization of the procedure encoding CDR3B and epitope amino acid sequences as numerical matrices. Each amino acid was mapped to a 6 dimensional row vector derived from published amino acid factor scores (36). CDR3B and epitope sequences of length less than 20 and 10, respectively, were padded to a predetermined length with placeholder amino acids (“-”), which were mapped to placeholder row vectors (Table S1). **(B)** Architecture of the hybrid neural network model consisting of the Triplet Network and Modal Alignment Network subunits. During training, all valid triplets (*x*_*pi*_, *x*_*ai*_, *x*_*ni*_) of CDR3B sequences were extracted from each minibatch of data and input into the Triplet Network, while the original minibatch (*x*_*i*_, *y*_*i*_) composed of binding CDR3B and epitope sequence pairs were input into the Modal Alignment Network. The Triplet and Modal Alignment Networks were trained simultaneously by combining their respective Triplet Loss and negative log likelihood loss functions into a single combined loss function. **(C)** Schematic of Triplet Network that trains the parameters of the CDR3B embedding network. The Triplet Network consisted of 3 copies of the CDR3B embedding network with shared parameters and took triplets of CDR3B sequences as input. **(D)** Schematic of Modal Alignment Network that trained the parameters of the epitope mapping network. The Modal Alignment Network took as input pairs of binding CDR3B and epitope sequences and calculated the negative log likelihood loss by taking the negative log of the probability of observing the latent space representation of the CDR3B sequence with respect to the Gaussian distribution obtained via epitope mapping network. **(E)** UMAP scatterplot visualization of CDR3B sequences embedded as points in latent space. Points were colored according to their binding epitope and shaped according to their membership in either the training or test set. Training the hybrid neural network model forced the CDR3B sequence embeddings to cluster in latent space by their binding epitope.

Additionally, we investigate several inconsistencies in the processing of CDR3B sequences by popular databases and the confounding effects of ignoring such inconsistencies on evaluating TCR-epitope binding prediction. The CDR3B sequence is a variable region of the TCR β chain flanked by conserved amino acids. A particular convention is usually followed when CDR3B sequences are reported in the literature; specifically, the N-terminus starts with a conserved cysteine (C) residue, while the C-terminus ends with a conserved phenylalanine (F) residue. CDR3B sequences that follow this convention are considered to be in “proper” form, while sequences not displaying this pattern are considered to be “improper.” Although some databases such as VDJDB (29) curate all CDR3B sequences into proper form, other databases such as IEDB (30) and McPAS (31) may contain a mixture of CDR3B in proper and improper forms. Importantly, we show that the preprocessing status may be correlated with distinct epitopes, thereby acting as a confounding factor and causing a machine learning model to learn the preprocessing status itself, rather than the intrinsic sequence patterns that confer TCR-epitope specificity.

In addition to the ability to predict TCR-epitope binding, it is also important to understand the features within CDR3B sequences determining epitope specificity. Previous studies have attempted to identify relevant CDR3B sequence motifs; however, these approaches are mostly limited to visualizing the empirical position-dependent amino acid distributions of CDR3B sequences binding a given epitope and do not describe features discriminating different epitopes. Interpreting the learned features of a machine learning model that can accurately predict TCR-epitope binding offers a powerful alternative strategy for extracting biologically relevant discriminatory features. In this work, we apply a statistical interpretation method, based on Markov Chain Monte Carlo (MCMC) sampling of input sequences (32,33), to extract the patterns of CDR3B sequences and biochemical properties that our model has learned to associate with specific epitope binding and to gain insight into the particular positions within the CDR3B domain important for binding. In contrast to the previous studies, such as (14), that have analyzed the physical structure of the TCR-pMHC complex first to constrain their motif analysis to the CDR3B regions contacting the epitope, our approach identifies the salient positions of CDR3B learned by our model without any prior structural information and then compares the results with known crystal structures of the TCR-pMHC complex. Our analysis shows that the inferred salient positions are not necessarily the closest to the epitope, highlighting a need to reconsider the common assumption that physical proximity represents importance for determining the TCR-epitope specificity. Our work thus advances key prediction tasks associated with TCR-epitope binding and provides interpretable insight into the learned features of our computational model.

## MATERIAL AND METHODS

### Positive binding and negative binding datasets

A dataset consisting of pairs of CDR3B sequences and their corresponding target epitope sequences, termed the positive binding dataset, was constructed by combining experimental data from four sources: IEDB (30), VDJdb (29), McPAS (31), and PIRD (34). The data were downloaded on February 26, 2020 and filtered as follows: removal of CDR3B sequences originating from TCRs of non-human hosts, restriction of data to pairs in which the epitope is presented by MHC class I molecules corresponding to HLA A*02:01, removal of pairs with either CDR3B or epitope sequence containing non-proteinogenic amino acids, removal of pairs with CDR3B sequence longer than 20 amino acids, and removal of pairs with epitope sequence longer than 10 amino acids. Constraints on CDR3B and epitope sequence lengths were imposed to accommodate the fixed-length representation used in our model, and the specific sequence lengths were chosen such that the vast majority of available CDR3B and epitope sequences were included in our analysis, while minimizing the input dimension. The sets of CDR3B-epitope pairs from the 4 databases were merged with duplicates removed. A set of CDR3B sequences assumed not to bind any epitopes represented in the positive binding dataset, termed the negative binding dataset, was constructed from the CDR3B sequences downloaded from (35) in August 2020, subjected to the following conditions: removal of sequences containing non-proteinogenic amino acids, removal of sequences with length greater than 20, retention of only those sequences conforming to the standard CDR3B conventions of having a C amino acid at the N-terminus and an F amino acid at the C-terminus, and removal of sequences represented within the positive binding dataset.

### CDR3B and epitope sequence representation

CDR3B and epitope amino acid sequences were both initially represented by strings of amino acid letters, as downloaded from the databases; however, our model required these sequences to be represented in a numeric form. Therefore, we formulated a numerical procedure to encode amino acid sequences into a specific matrix form which we call the Atchley representation. We first mapped each amino acid to a 6-dimensional vector, which we termed the Atchley amino acid vector, capturing various physical and biochemical properties of the amino acid according to a factor analysis of 54 features (36). More specifically, the first 5 entries of the 6-dimensional vector representation of an amino acid corresponded to the 5 factor scores associated with the amino acid transformed by min-max scaling, and the remaining entry was set to 1 as an indicator of being a real amino acid. Supplementary Table S1 displays the vector representations of the amino acids. This encoding scheme allowed us to represent a sequence of amino acids as a matrix with the 6-dimensional vectors along rows. Furthermore, to accommodate CDR3B or epitope sequences of varying lengths, we padded the matrix representation of each CDR3B and epitope sequence with placeholder row vectors to achieve a total row dimension of 20 and 10, respectively. The placeholder row vectors contained a value of 0.5 for the first 5 entries and 0 in the final entry. We therefore mapped CDR3B and epitope amino acid sequences to Atchley amino acid matrices of dimension 20 × 6 and 10 × 6, respectively (Figure 1A).

### Overview of model architecture

Our model consists of two neural networks that are used to calculate the unnormalized binding affinity between input TCR and epitope. The first neural network, termed the CDR3B embedding network, takes an input Atchley representation (Methods: CDR3B and epitope sequence representation) of a TCR CDR3B sequence *x* and outputs a 32-dimensional vector *f*(*x*). The second neural network, termed the epitope mapping network, takes an input Atchley representation of an epitope sequence *y* and outputs two 32-dimensional vectors, *μ*(*y*) and *σ*(*y*). A diagonal matrix Σ(*y*) is then constructed with main diagonal entries taken from the vector *σ*(*y*). The unnormalized binding affinity *ρ* between the TCR and epitope is then calculated as follows:

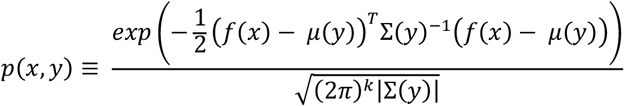

where |Σ(*y*)| is the determinant of Σ(*y*).

Intuitively, the CDR3B embedding network maps the Atchley representation *x* of a CDR3B sequence to point *f*(*x*) in ℝ^32^, while the epitope mapping network maps the Atchley representation *y* of an epitope sequence to the parameters *μ*(*y*) and *σ*(*y*) of a Gaussian distribution with diagonal covariance matrix and support over the same latent space ℝ^32^. The unnormalized binding affinity is defined to be the probability of observing the CDR3B sequence represented by *f*(*x*) with respect to the Gaussian distribution of the epitope sequence parametrized by *μ*(*y*) and *σ*(*y*).

To enable our model to predict TCR-epitope binding, we tuned the parameters of both the CDR3B embedding network and epitope mapping network with two goals in mind: first, the CDR3B embedding network must map the Atchley representations of CDR3B sequences to points in the ℝ^32^ latent space such that the CDR3B sequences of TCRs binding the same epitope are clustered together into the same point cloud in latent space; second, the epitope mapping network must map the Atchley representations of epitope sequences to Gaussian distributions over the latent space, localized to the point clouds of their corresponding CDR3B sequences. To accomplish the first goal, we constructed a Triplet Network, composed of three copies of the CDR3B embedding network trained using a Triplet Loss function (Supplementary Methods: Model architecture and training). The input to the Triplet Network was a set of triplets of CDR3B Atchley representations. For each triplet in the input set, two of the CDR3B Atchley representations, known as the anchor and the positive, corresponded to TCRs binding the same epitope, while the third remaining CDR3B Atchley representation, known as the negative, was derived from the negative binding set or corresponded to a TCR binding a different epitope. Training the Triplet Network with Triplet Loss was designed to tune the parameters of the CDR3B embedding network to cluster latent space representations of CDR3B sequences according to their corresponding binding epitope by pulling together the latent space representations of the anchor and positive and pushing apart the latent space representations of the anchor and negative for every triplet in the input triplet set (Supplementary Methods: Model architecture and training). To accomplish the second goal, we constructed a network termed the Modal Alignment Network. The Modal Alignment Network took as input the Atchley representations of a pair of binding CDR3B and epitope sequences – denoted by *x* and *y*, respectively – and was trained to minimize the negative log likelihood loss function −log (*p*(*x*, *y*)). Given that *p*(*x*, *y*) is the unnormalized binding affinity, minimizing the loss function of the Modal Alignment Network tuned the parameters of the epitope mapping network to map the Atchley representations of epitopes to Gaussian distributions that maximized the likelihood of observing the latent space representations of their corresponding binding CDR3B sequences. During the model training process, both the Triplet and Modal Alignment Networks were trained simultaneously by minimizing a combined loss function comprising of Triplet Loss and negative log likelihood loss (Supplementary Methods: Model architecture and training). The weights of the CDR3B embedding network components contained in the Triplet and Modal Alignment Networks were shared to ensure that the CDR3B embedding network accomplished both goals of clustering the representations of CDR3B sequences into point clouds in latent space according to their binding epitope and aligning each point cloud to the Gaussian distribution corresponding to its binding epitope.

### Model training

We analyzed the performance of our model on Tasks 1 and 2 using various subsets of the entire dataset with varying number and composition of epitope classes in the training and test sets. Each subset was specified by a set of epitopes, called the “seen epitope set,” and included their bound TCRs. For each given subset, we trained the above model with a single training procedure and used a single test set to evaluate Tasks 1 and 2 by performing two different evaluation procedures. In order to build the training and test sets for a given seen epitope set, we placed 80% of the CDR3B sequences from the positive binding set (Methods: Positive binding and negative binding datasets), corresponding to TCRs binding an epitope in the seen epitope set, into the training set, while the remaining 20% were placed into the test set. The size and composition of the training and test sets could therefore be specified by the number and identity of the epitopes in the seen epitope set. Since the number of examples associated with each epitope class is highly imbalanced (Supplementary Table S2), we must balance the classes to ensure that our model learns to predict TCR-epitope binding for minority classes. Therefore, during formation of training and test sets, we upsampled the minority classes with replacement until all classes contained the same number of examples as the majority class. Training and test sets were upsampled independently to ensure that no single CDR3B sequence appeared in both the training and test sets simultaneously. Given *N*, the size of the training set after upsampling, 0.5 × *N* CDR3B sequences were randomly sampled without replacement from the negative binding set (Methods: Positive binding and negative binding datasets) and added to the training set. Additionally, 10,000 CDR3B sequences were randomly sampled without replacement from the remaining negative set and added to the test set.

Training was performed by first randomly shuffling the training dataset for every epoch and utilizing minibatches of size 128. For each minibatch, all valid triplets of Atchley representations of CDR3B sequences were extracted and input into the Triplet Network. Additionally, the pairs of Atchley representations of CDR3B sequences and epitope sequences contained in the minibatch were input into the Modal Alignment Network. Both the Triplet Network and Modal Alignment Network were trained simultaneously by minimizing an overall loss function that combined the Triplet Loss and negative log likelihood (Supplementary Methods: Model architecture and training) and using early stopping for a set number of epochs determined by the seen epitope set.

### Definition of classification tasks and general model evaluation procedure

Task 1 was defined as follows: given a predetermined epitope sequence and TCR repertoire, classify the TCRs within the given TCR repertoire into binding and non-binding classes. To evaluate the model on Task 1, a query epitope *y* was first chosen from the seen epitope set. A query CDR3B sequence set corresponding to the query epitope was then constructed from the test set by retaining only the CDR3B sequences corresponding to TCRs binding to the query epitope as well as all CDR3B sequences derived from the negative set. The unnormalized binding affinity between the query epitope and every CDR3B sequence in the test set was then calculated. The Area Under the ROC Curve (AUC) metric was then measured from the unnormalized binding affinity scores. This procedure was repeated with each epitope in the seen epitope set taking on the role of the query epitope.

Task 2 was defined as follows: given a set of TCRs known to bind one of the epitopes in the seen epitope set, identify the binding epitope for each TCR. To construct the query CDR3B sequence set specific to Task 2, the CDR3B sequences from the negative set were first removed from the test set. For each CDR3B sequence in the query CDR3B sequence set, its associated unnormalized binding affinity to each epitope in the epitope set was calculated. The TCR corresponding to the CDR3B sequence was then predicted to bind the epitope corresponding to the highest unnormalized binding probability. Classification accuracy was calculated as the percentage of all TCRs with CDR3B sequence contained in the test set predicted to bind their true binding epitopes.

### Processing of CDR3B sequences

CDR3B sequences displaying conserved C and F residues at the N- and C-terminus, respectively, were termed as “proper CDR3B” sequences in this manuscript. CDR3B sequences not curated in the databases with both of these conserved residues were similarly termed as “improper CDR3B” sequences. We sought to process improper CDR3B sequences into proper form and denoted the converted sequences as “fixed proper CDR3B” sequences. CDR3B sequences already in proper form without the application of this correction procedure were termed “native proper CDR3B” sequences. In the procedure, we first counted the occurrences of 3-mers following the conserved C residue at the N-terminus of native proper CDR3B sequences and the occurrences of 3-mers at the N-terminus of improper CDR3B sequences without the conserved C residue (Supplementary Table S3). Similarly, we counted the occurrences of 3-mers directly preceding the conserved F residue at the C-terminus of native proper CDR3B sequences and the occurrences of 3-mers at the C-terminus of improper CDR3B sequences without the conserved F residue (Supplementary Table S3). An improper CDR3B sequence lacking a conserved C residue at the N-terminus was considered to have a fixable-at-*k* N-terminus if its N-terminal 3-mer appeared within the top *k* occurring N-terminal 3-mers of both proper and improper CDR3B sequences. Improper CDR3B sequences with a fixable-at-*k* N-terminus could have their N-terminus fixed into proper form by appending the C residue prefix. Likewise, an improper CDR3B sequence lacking a conserved F residue at the C-terminus was considered to have a fixable-at-*k* C-terminus if its C-terminal 3-mer appeared within the top *k* occurring C-terminal 3-mers of both the proper and improper CDR3B sequences or contained the XFG motif, with X representing an arbitrary amino acid. CDR3B sequences with a fixable-at-*k* C-terminus could have their C-terminus processed into proper form by removing the extraneous G residue when the C-terminal 3-mer contained the XFG motif or by appending the F residue suffix. Improper CDR3B sequences that could be processed into proper form by fixing N- and/or C-termini at a given value of *k* were considered to be “fixable improper CDR3B” sequences. Improper CDR3B sequences that could not be processed at a given value of *k* were considered “unfixable improper CDR3B” sequences. We thus created the “unfixed dataset,” consisting of native proper and fixable improper CDR3B sequences, and the “fixed dataset,” consisting of native proper and fixed proper CDR3B sequences, by performing the above procedure on CDR3B sequences in the positive binding set with *k* set to 10. The value of *k* was chosen to include only the most highly represented 3-mers.

### Identification of preprocessing artifacts as confounding factors

The model was trained with training and test sets constructed from the unfixed dataset with the seen epitope set specified to include only epitopes GILGFVFTL and NLVPMVATV. The training was performed using early stopping at 250 epochs. To investigate the consequences of ignoring the data preprocessing steps on model performance for Task 2, we explored the relationship between the presence and absence of the conserved C residue at the N-terminus of CDR3B sequences and the classification status of the CDR3B sequence as corresponding to a TCR binding either GILGFVFTL or NLVPMVATV. Given the Atchley representations *y*_*GILGFVFTL*_ and *y*_*NLVPMVATV*_ of the epitope sequences GILGFVFTL and NLVPMVATV, respectively, we calculated the log ratio of the unnormalized binding affinities

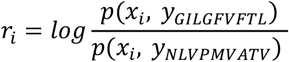

for the Atchley representation of each CDR3B sequence in the test set, denoted by *x*_0_, under two settings: with the CDR3B sequence displaying proper N- and C-terminus and with the CDR3B sequence truncated to remove the conserved C residue at the N-terminus while still displaying proper C-terminus. We performed a two-sided Mann-Whitney U test to measure the divergence of the spreads of *r*_*i*_ under the proper N-terminus and truncated N-terminus settings.

To investigate the effects of the presence or absence of processing of CDR3B sequences on classification accuracy in the Task 2 classification setting, the model was trained using early stopping at 250 epochs and 5-fold cross-validation with 80-20 train-test splits of the CDR3B sequences of TCRs binding the epitopes GILGFVFTL and NLVPMVATV. Classification accuracy of the test sets was evaluated in 4 different contexts (Results: Artifacts in preprocessing CDR3B sequences confound TCR-epitope binding predictions).

### Model evaluation for specific seen epitope sets

We first ranked the epitopes according to their number of annotated binding TCR CDR3B sequences in the fixed dataset (Supplementary Table S2). We then trained and evaluated the model on Tasks 1 and 2 using 5-fold cross-validation on 4 pairs of training and test sets derived from the fixed dataset. Each pair of training and test sets was constructed by taking the top *M* epitopes as the seen epitope set, with *M* taking values between 2 and 5. For *M* = 2, 3, 4, and 5, training was performed using early stopping at 250, 180, 140, and 120 epochs, respectively. The number of epochs was chosen by training the model for 500 epochs and determining the epoch at which test loss started to rise. To investigate the performance of the model in the setting with a greater number of classes and fewer examples per class, we created the “reduced fixed dataset” by removing CDR3B sequences binding the top 5 epitopes from the fixed dataset. We then repeated the evaluation procedure on 4 pairs of training and test sets derived from the reduced fixed dataset, with each pair of sets specified by taking the top *P* epitopes of the reduced fixed dataset as the seen epitope set and *P* taking on values of 5, 10, 15, and 20. Training was performed using early stopping at 500 epochs for all values of *P*.

### Model interpretation

We first trained our model on the training set specified by taking the top 4 epitopes as the seen epitope set. We then interpreted the learned features of our model using an MCMC method inspired by (32,33), fixing the epitope sequence input to the model and sampling the space of CDR3B sequences of fixed length with a bias towards CDR3B sequences with higher unnormalized binding affinity for the specified epitope (Supplementary Methods: MCMC method of model interpretation). Since our MCMC method varies the inner positions of the CDR3B sequence while fixing the epitope sequence and CDR3B sequence length, our method interprets the features that the model has learned to predict binding between a specific epitope and CDR3B sequences of a specific length. The MCMC method was performed for 12 combinations of CDR3B sequence length and epitope sequence, corresponding to CDR3B sequences of lengths 13, 14, and 15 associated with TCRs binding the epitopes GILGFVFTL, NLVPMVATV, GLCTLVAML, and ELAGIGILTV. For each combination of CDR3B sequence length and epitope sequence, the CDR3B sequences associated with TCRs binding the given epitope were selected as candidates to seed the MCMC runs. From this candidate set, the top 40 CDR3B sequences with the highest predicted binding affinities from the training and test set were selected as seeds for the MCMC runs. Five MCMC runs were initiated at each seed, with each run proceeding for 20,000 steps, resulting in a total of 200 MCMC runs, termed the “individual MCMC runs,” for each combination of CDR3B sequence length and epitope sequence. Inverse temperature parameter *β* was set to 2.8, 2.8, 2.0, and 1.6 for MCMC runs associated with the epitopes GILGFVFTL, NLVPMVATV, GLCTLVAML, and ELAGIGILTV, respectively. MCMC runs sharing epitope sequence and input CDR3B length were clustered by agglomerative clustering utilizing complete linkage, with the distance metric between runs defined as the Jenson-Shannon Divergence (JSD) between their associated empirical amino acid distributions. The number *Q* of clusters was varied between 2 and 40, and the Silhouette score was measured for each value of *Q*. For values of *Q* displaying the highest peaks in the Silhouette score, clustered MCMC runs were combined, and sequence logos were generated from the empirical amino acid distributions derived from the sampled CDR3B sequences using the Python package weblogo (37). Each combined MCMC run corresponding to a cluster, termed the “cluster MCMC run,” was then ranked according to a uniqueness score constructed as follows: first, for each combination of CDR3B sequence length and epitope sequence, all 200 MCMC runs were combined into a single MCMC run, termed the “representative MCMC run”; next, every cluster MCMC run for a given CDR3B sequence length and epitope sequence was given a uniqueness score calculated by averaging its JSD with all representative MCMC runs associated with the same CDR3B sequence length and alternative epitope sequence.

To investigate the physical and biochemical properties of amino acids found to affect the binding affinity of CDR3B sequences, we constructed a 5 × *L* matrix from each individual MCMC run, with the 5 rows corresponding to the 5 entries of the amino acid vector representation (36) and the *L* columns corresponding to the middle *L* positions in the CDR3B sequence. For each position in the CDR3B sequence, a weighted average of amino acid vectors was taken over the MCMC samples, with the weight of each amino acid vector given by the empirical frequency of its corresponding amino acid at that position. Matrices constructed from individual MCMC runs in this manner were termed “individual matrices.” For each combination of CDR3B sequence length and epitope sequence, we also generated a single matrix, termed the “representative matrix,” by performing the same procedure on the corresponding representative MCMC run. Additionally, for each combination of CDR3B sequence length and epitope sequence, we clustered the individual matrices via agglomerative clustering utilizing complete linkage, with the distance metric between matrices defined as the Frobenius distance. The number of clusters *R* was varied between 2 and 40, and the Silhouette score was measured for each value of *R*. For each combination of CDR3B sequence length and epitope sequence, the final number of clusters was chosen to be the value of *R* displaying the highest peak in Silhouette score. After the number of clusters was determined, each cluster of matrices was associated with a single matrix, termed the “cluster matrix,” constructed by averaging the matrices within the cluster. The resulting cluster matrices were then ranked according to a uniqueness score given by its average Frobenius distance to representative matrices with identical CDR3B sequence length and alternative epitope sequence. A more detailed description of the MCMC method and clustering is given in Supplementary Methods: MCMC method of model interpretation.

### TCR-pMHC structure analysis

To investigate where contact between the CDR3B region of the TCR and the epitope occurs, a set of TCR-pMHC structures was obtained from the database STCRDab (38). We filtered the set for structures with structure resolution finer than 3 Angstroms and in which one of the top 4 epitopes (GILGFVFTL, NLVPMVATV, GLCTLVAML, ELAGIGILTV) was presented on a class I MHC molecule, resulting in 17 structures. The resolution threshold was chosen to remove data with high uncertainty in atomic and residue positions (39). For each structure, the pairwise distance between each epitope amino acid and CDR3B region amino acid was computed, yielding a 2-dimensional distance matrix. The distance matrices of structures containing the same epitope binding to CDR3B sequences of the same length were aggregated using an elementwise average. The MCMC method of model interpretation was run on the same combinations of CDR3B sequence length and epitope sequence as were present in the TCR-pMHC structural data, with the inverse temperature parameter *β* set to 1.7, 1.7, 1.7, and 1.3 for the epitopes GILGFVFTL, NLVPMVATV, GLCTLVAML, and ELAGIGILTV, respectively. We sought to measure the correlation between the median distance of a position on the CDR3B region to the epitope, as given by structural data, and the salience of a position on the CDR3B region (Supplementary Methods: MCMC method of model interpretation). Due to the high conservation of the CDR3B sequence at positions near the C-terminus and N-terminus, the first and last 4 amino acid positions of the CDR3B were excluded, retaining only the highly variable center region of the CDR3B for analysis. For each of the remaining positions in the CDR3B, the median distance to the amino acids of the epitope was calculated. Each of the 17 structures was matched with the set of MCMC runs corresponding to the same combination of epitope and CDR3B length. For each combination, the position-wise KL divergence was calculated from the sampled CDR3B sequences derived from the MCMC run. The 4 leading and 4 trailing positions were excluded, and the remaining positions corresponding to the highly variable CDR3B region were ranked by KL divergence in decreasing order, with the position with highest KL divergence receiving a rank of 0. For each position in the CDR3B region, its rank was summed over the MCMC runs and standardized yielding the “salience rank.” For each position in the CDR3B region in each of the 17 structures, the median distance and salience rank was plotted, and the Pearson correlation coefficient and p-value were calculated.

## RESULTS

### A hybrid neural network represents CDR3B and epitope sequences in a shared latent space

We designed a hybrid neural network model that can simultaneously perform Task 1 and Task 2 (Methods: Definition of classification tasks and general model evaluation procedure) by learning representations of both CDR3B and epitope sequences in a shared latent space, given an initial procedure encoding amino acid sequences into numerical matrices that embody various physical and biochemical properties (36) (Figure 1A; Methods: CDR3B and epitope sequence representation). More specifically, our model consisted of two main components (Figure 1B): a Triplet Network (25,26) that trained a CDR3B embedding network to embed CDR3B sequences into a latent space, clustering them into point clouds according to their binding epitope, and a Modal Alignment Network that trained an epitope mapping network to represent epitopes as Gaussian distributions over latent space localized to their corresponding CDR3B point clouds (Figure 1E). The Triplet and Modal Alignment Networks themselves consisted of CDR3B embedding network and epitope mapping network subunits (Figure 1C, Supplementary Figure S3 for Triplet Network and Figure 1D, Supplementary Figure S4 for Modal Alignment Network; Methods: Overview of model architecture, Supplementary Methods: Model architecture and training). The architecture and weights of the CDR3B embedding network subunits were shared within the Triplet Network and also between the Triplet Network and the Modal Alignment Network, which were trained together (Methods: Model training).

### Artifacts in preprocessing CDR3B sequences confound TCR-epitope binding predictions

Previous studies, such as NetTCR (21) and ERGO (23), attempting to predict the binding epitopes of TCRs from their CDR3B sequences have not explicitly addressed the potential nonuniform preprocessing of CDR3B sequences in public databases. Training and testing computational models on CDR3B datasets not standardized via a proper preprocessing step (Methods: Processing of CDR3B sequences) may inadvertently force the models to learn epitope-specific artifacts and fictitiously inflate the assessment of model performance. CDR3B sequences listed in some databases are preprocessed into proper CDR3B sequences which display conserved C and F amino acids at the N- and C-terminus, respectively. Other databases of TCR-epitope binding interactions, however, may contain a mixture of “proper CDR3B” sequences and “improper CDR3B” sequences (Methods: Processing of CDR3B sequences). Common forms of improper CDR3B sequences include sequences missing the single conserved C or F amino acid at the N- or C-terminus, respectively, as well as sequences retaining the superfluous conserved FGXG motif at the C-terminus, with X representing an arbitrary amino acid. Furthermore, the preprocessing status of a CDR3B sequence might provide information on the identity of its binding epitope, as the ratio of the number of reported proper CDR3B sequences to the number of reported improper CDR3B sequences varies widely between epitopes (Figure 2A; Supplementary Figure S5).

**Figure 2:**
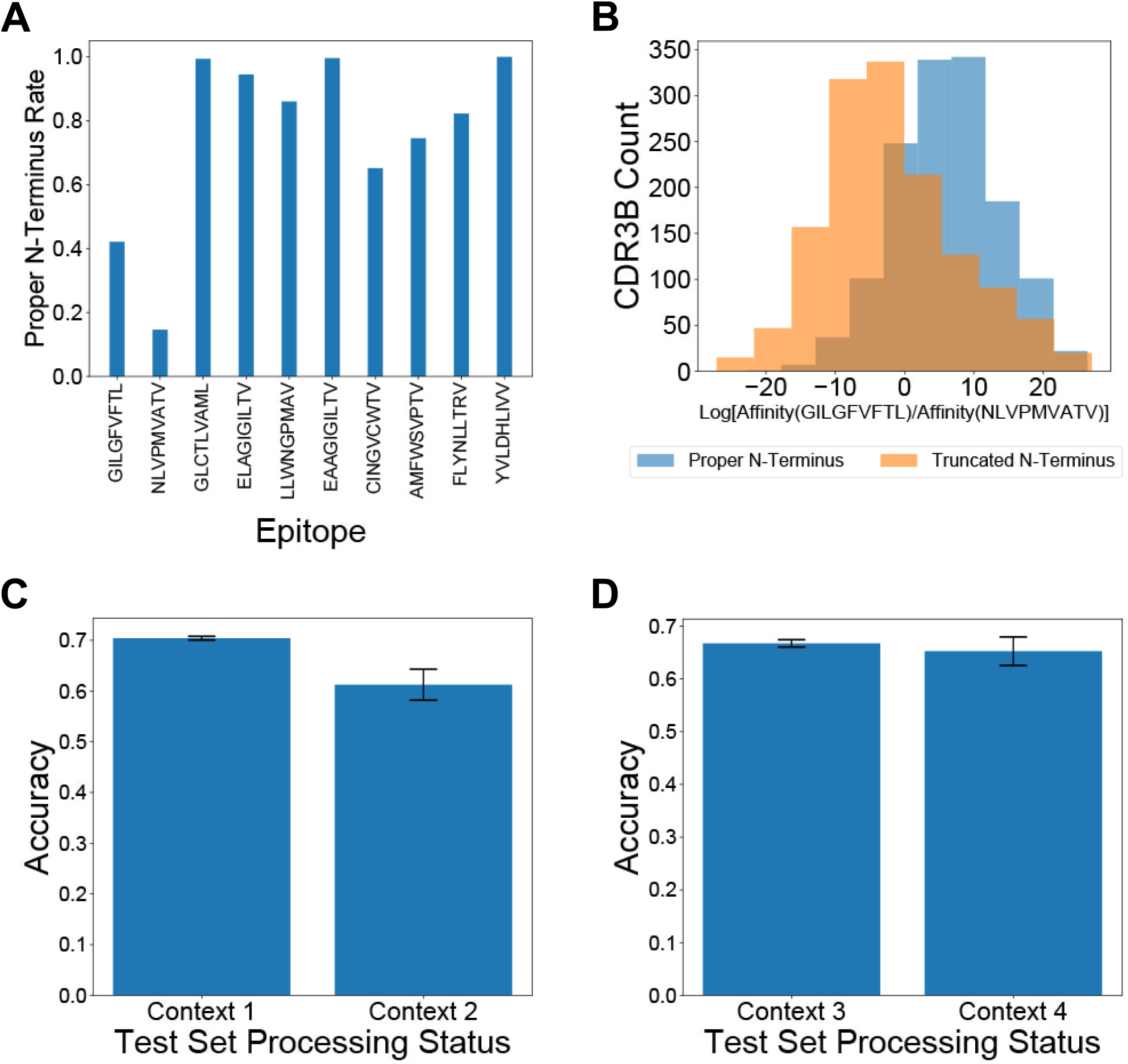
Processing CDR3B sequences reduces bias in Task 2 classification. **(A)** Proportion of CDR3B sequences derived from the unfixed dataset displaying proper N-terminus, arranged by binding epitope. **(B)** Histograms displaying the distribution of log ratios of unnormalized binding affinity towards GILGFVFTL versus NLVPMVATV for CDR3B sequences in the test set. Sequences with proper N-terminus showed a bias towards positive values, associated with preferential classification as binding the epitope GILGFVFTL, demonstrating that the model had learned epitope-specific processing artifacts. **(C)** Classification accuracy for CDR3B sequences binding GILGFVFTL or NLVPMVATV under Context 1 and Context 2. Discrepancy in classification accuracy between Context 1 and Context 2 demonstrated that ignoring preprocessing of CDR3B sequences may inflate assessment of the model on TCR-epitope binding prediction. **(D)** Classification accuracy for CDR3B sequences binding to GILGFVFTL and NLVPMVATV under Contexts 3 and 4. The similarity of classification accuracy between Context 3 and Context 4 demonstrated that preprocessing of CDR3B sequences did not introduce any new biases that would affect model assessment.

Without uniform processing of CDR3B sequences, machine learning models might learn to identify the binding epitope of CDR3B sequences based on the processing status of the N- and C-termini, rather than the inherent patterns determining epitope binding. For example, over 40% of CDR3B sequences binding GILGFVFTL display a proper N-terminus in the unfixed dataset (Methods: Processing of CDR3B sequences), while less than 20% of CDR3B sequences binding NLVPMVATV display a proper N-terminus (Figure 2A; Methods: Processing of CDR3B sequences). Therefore, a model trained on the unfixed dataset might assign a higher probability of binding GILGFVFTL to a CDR3B sequence containing a proper N-terminus relative to the variant of the same sequence containing an improper N-terminus. To demonstrate this effect, we trained our model on 80% of the CDR3B sequences corresponding to TCRs binding the epitopes GILGFVFTL and NLVPMAVTV in the unfixed dataset, containing a mixture of proper and improper CDR3B sequences (Methods: Processing of CDR3B sequences, Model training). We evaluated the log ratio of unnormalized binding affinity to GILGFVFTL versus unnormalized binding affinity to NLVPMVATV for the remaining 20% of CDR3B sequences under two processing conditions: with all CDR3B sequences processed to have the conserved C and F residues at their respective N- and C-terminus, and with the same CDR3B sequences excluding the conserved C residue at the N-terminus while retaining the conserved F residue at the C-terminus (Methods: Identification of preprocessing artifacts as confounding factors). The distributions of log ratios of unnormalized binding affinity showed a clear bias of the proper CDR3B sequences towards being classified as binding the epitope GILGFVFTL (Figure 2B; *p* = 1.83×10^−116^, two-sided Mann-Whitney U test), confirming that the model indeed learned the epitope-specific processing artifacts to perform classification.

To understand the effect of including unprocessed CDR3B sequences on evaluation of Task 2 (Methods: Definition of classification tasks and general model evaluation procedure), we classified CDR3B sequences as binding GILGFVFTL versus NLVPMVATV using 5-fold cross-validation under four different contexts (Methods: Identification of preprocessing artifacts as confounding factors): both training and test sets derived from the unfixed dataset (Context 1); unfixed training set, but test set filtered to include only native proper CDR3B sequences (Context 2); both training and test sets derived from the fixed dataset (Context 3); and, fixed training set, but the test set filtered to include only native proper CDR3B sequences (Context 4). Context 1 simulated the setting in which researchers failed to process the CDR3B sequences in both training and test sets. In this context, a relatively high test accuracy may arise from the epitope-specific processing status artificially informing the model about the associated epitope of a CDR3B sequence. Context 2 simulated the setting in which researchers trained their model on unprocessed CDR3B sequences and proceeded to test their model on an external set of proper CDR3B sequences. As expected, the classification accuracy in Context 2 was diminished compared to Context 1, because the processing status no longer provided any epitope-specific information for the test CDR3B sequences (Figure 2C), suggesting that the model performance was artificially over-estimated in Context 1. Context 3 simulated the condition in which researchers processed all CDR3B sequences into proper form prior to training and testing. Context 4 simulated the setting in which researchers trained their model on processed CDR3B sequences and proceeded to test their model on an external set of proper CDR3B sequences. The classification accuracy in Context 4 was higher than that in Context 2 (Figure 2C, D), demonstrating that training on properly processed data successfully removed the confounding factor of the preprocessing status and improved the model performance on unseen proper CDR3B sequences. Furthermore, the classification accuracies were similar between Context 3 and Context 4 (Figure 2D), supporting that our processing procedure did not introduce any new biases.

These results demonstrated the critical importance of uniform preprocessing of CDR3B sequences prior to model training and evaluation for accurate assessment of model performance and generalizability. We therefore utilized the fixed dataset for all subsequent evaluation and analysis of our model.

### The trained model predicts TCR-epitope binding under Task 1 and Task 2 settings

We first evaluated the model on Task 2 of assigning CDR3B sequences to epitope classes containing the most amount of data in the fixed dataset (Methods: Processing of CDR3B sequences), with the number *M* of epitope classes varying from 2 to 5 (Methods: Model evaluation for specific seen epitope sets). We computed the cross-validation classification accuracy for the different values of *M* and observed a decrease in classification accuracy for increasing values of the class number (Figure 3A), as to be expected for multi-class classification tasks. The classification accuracy was, however, greater than random for all values of *M*, indicating that the model was learning to identify the binding epitope of TCRs from their CDR3B sequence. For each value of *M*, we also evaluated the mean performance of the model on Task 1 of predicting whether the CDR3B sequences could bind each given epitope (Figure 3B, Supplementary Table S4). The mean AUC scores ranged from 0.66 to 0.80, demonstrating that the model was learning to perform the binary classification task.

**Figure 3:**
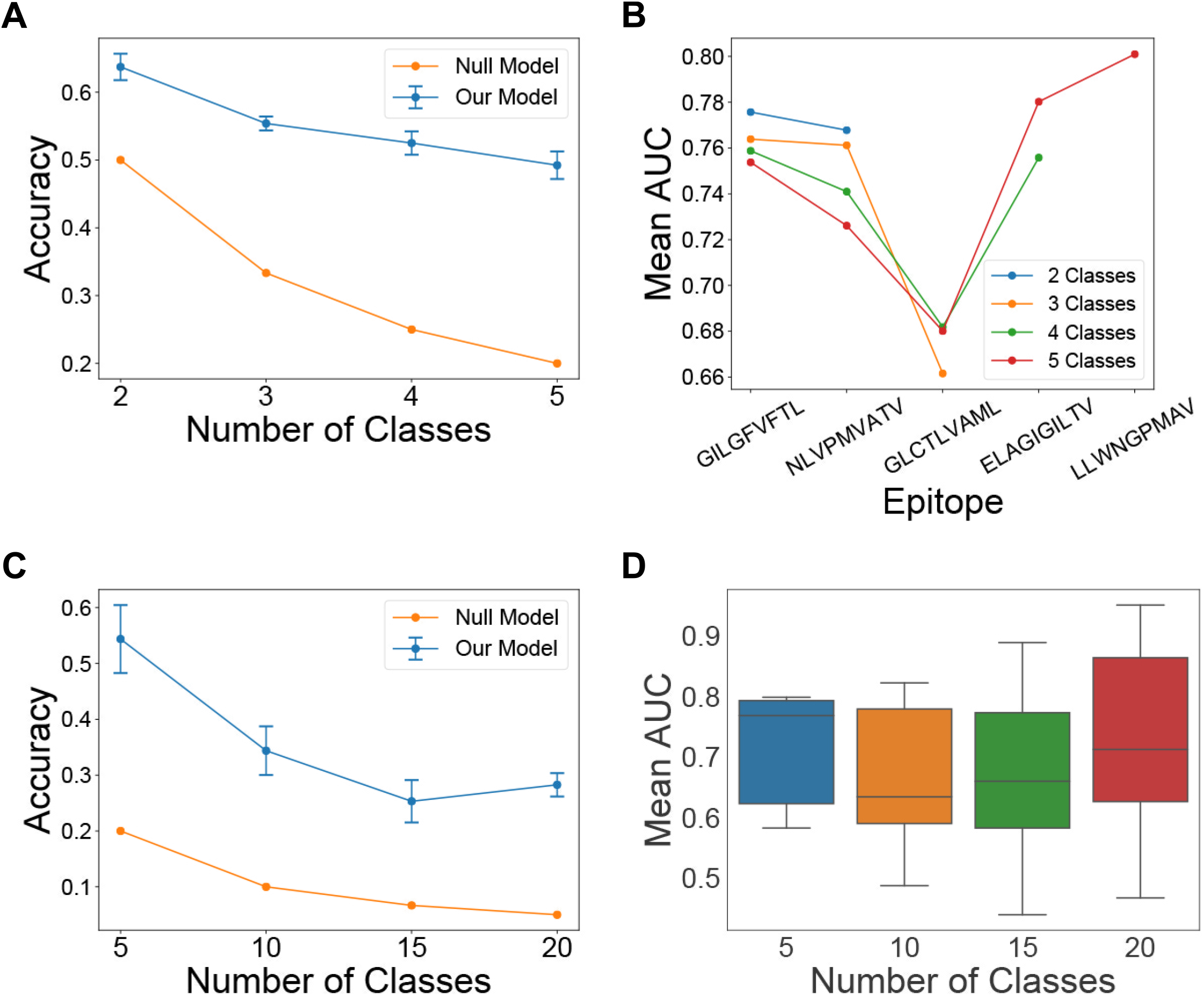
Model performance on multiclass and binary classification tasks. **(A)** Line chart displaying the mean and standard deviation of the 5-fold cross-validation classification accuracy of the model on Task 2 using the fixed dataset, with the number of epitope classes shown on the *x*-axis. Classification accuracy of a null model performing random classification is also shown for reference. **(B)** Line chart displaying the mean AUC score of the model on Task 1 for each epitope class displayed on the *x*-axis, with the number of classes varying from 2 to 5. A separate line is plotted for each value of the number of classes **(C)** Line chart displaying the mean and standard deviation of the 5-fold cross-validation classification accuracy of the model on Task 2 for the reduced fixed dataset, with the number of classes shown on the *x*-axis. Classification accuracy of a null model performing random classification is also shown for reference. **(D)** Box plot of the distribution of AUC scores for the model on Task 1 using the reduced fixed dataset, with the number of epitope classes shown on the *x*-axis.

Since the vast majority of epitope sequences represented in the fixed dataset have very few binding CDR3B sequences, we also sought to evaluate the performance of the model on Tasks 1 and 2 under the condition of limited data. We first constructed a “reduced fixed dataset” (Methods: Model evaluation for specific seen epitope sets) from the fixed dataset by excluding all CDR3B sequences binding one of the top 5 epitopes with the most recorded binding CDR3B sequences (Supplementary Table S2). We then evaluated the model on the top *P* epitope classes with the most data in the reduced dataset, with the number *P* of epitope classes set to 5, 10, 15, and 20 (Figure 3C for Task 2 and Figure 3D for Task 1; Methods: Model evaluation for specific seen epitope sets). The model was able to perform Task 2 with classification accuracy greater than random chance (Figure 3C), suggesting that the model could perform Task 2 even with very little data and a large number of classes. However, the model showed wide variability in performance on Task 1 under this setting (Figure 3D), with mean AUC dropping below 0.5 for certain classes, suggesting that data availability is critical for performing Task 1.

To assess our model with respect to other state-of-the-art approaches, we compared the performance of our model on Task 1 to Table 2 from Springer et al. (23) summarizing the performance of their model on Task 1 relative to the performance of a model based on Gaussian Processes (20). The authors evaluated two different versions of their model on 3 different datasets: McPAS, VDJdb, and a combination of McPAS and VDJdb. We analyzed the dataset consisting of a combination of McPAS and VDJdb datasets, as it most closely reflected the setting in which we evaluated our model. Of note, the classification task we performed differed from that performed by Springer et al. Our model was trained to identify for a specified epitope its binding TCRs from a set consisting of TCRs that bind the given epitope and ambient TCRs, derived from a healthy control group of donors and not contained in the positive binding set (Methods: Positive binding and negative binding datasets). By contrast, Springer et al. performed a classification task, which they termed single peptide binding (SPB), using a set consisting of TCRs binding the specified epitope and TCRs known to bind alternative epitopes. Our model had the best performance on Task 1 with respect to the epitope NLVPMVATV and showed competitive performance with respect to the epitopes GILGFVFTL and GLCTLVAML (Supplementary Table S6).

### Model interpretation method identifies salient CDR3B sequence motifs and amino acid positions specifying epitope binding

We sought to discover the CDR3B sequence features that determine binding affinity to the top 4 epitopes with the most data (GILGFVFTL, NLVPMVATV, GLCTLVAML, and ELAGIGILTV), using a neural network interpretation method based on the maximum entropy principle (32) (Methods: Model Interpretation and Supplementary Methods: MCMC method of model interpretation). For each given epitope, this probabilistic interpretation method performed successive single amino acid mutagenesis starting from a seed CDR3B sequence known to bind the epitope, with the MCMC sampling biased towards sequences with high predicted affinity and away from sequences with low predicted affinity. We summarized the MCMC runs by visualizing the empirical amino acid distribution at each position of the CDR3B sequence across the MCMC samples using weblogos (37). Positions within the CDR3B sequence permitting a wide variety of residues with minimal impact to predicted binding affinity displayed low sequence conservation across the MCMC samples, while positions at which specific amino acids were required for high predicted binding affinity were preserved during sampling. To highlight the advantage of the MCMC interpretation method over simply aligning the CDR3B sequences reported in the dataset, we first visualized the weblogos of the aligned CDR3B sequences of length 15 binding the top 4 epitopes (Figure 4A). This visualization method simply extracted a conserved amino acid motif from the empirical distribution of CDR3B sequences binding an epitope and had difficulty discovering individual motifs determining the specificity of CDR3B-epitope binding. For example, the CDR3B sequences for all 4 epitopes exhibited a conserved CASS motif at the C-terminus (positions 1-4), a conserved EQYF motif at the N-terminus (positions 12-15), and conserved glycine residues at positions 7-9 (Figure 4A). These shared features thus could not discriminate specific interactions between epitopes and TCRs.

**Figure 4:**
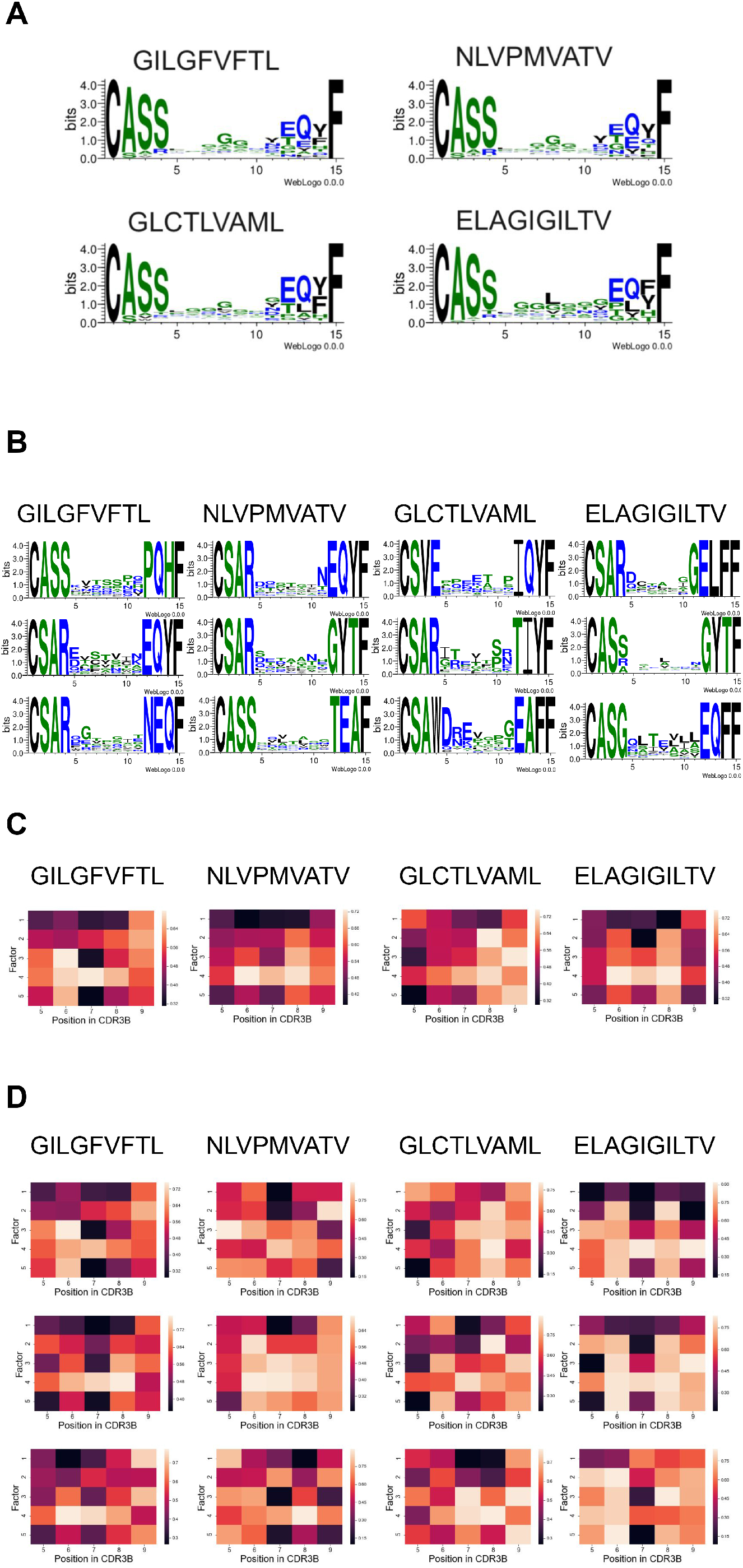
Consensus and discriminative CDR3B motifs and their associated biochemical properties extracted by the MCMC interpretation method. **(A)** Weblogos visualizing the alignment of CDR3B sequences of length 15 binding the top 4 epitopes. **(B)** Weblogos corresponding to the top 3 most unique cluster MCMC runs for CDR3B sequences of length 15 binding the top 4 epitopes. **(C)** Heatmap visualization of the representative matrices of CDR3B sequences of length 13 binding the top 4 epitopes. Each column indexes a position in the CDR3B sequence, and each row corresponds to one of 5 factor scores associated with amino acid chemical properties. **(D)** Heatmap visualization of the top 3 most unique cluster matrices for CDR3B sequences of length 13 binding the top 4 epitopes.

We simulated 200 MCMC chains for each pair of CDR3B sequence length and epitope sequence, with the seeds for each pair chosen to be the top 40 CDR3B sequences of the specified length with the highest unnormalized binding affinity to the specified epitope (Methods: Model interpretation) and 5 independent MCMC runs initiated from each seed. MCMC runs constructed in this manner were termed the “individual MCMC runs.” In contrast to the simple alignment method, the MCMC method, which interpreted the learned features of our model that was designed to discriminate the CDR3B sequences of TCRs binding different epitopes, highlighted epitope-specific CDR3B sequence motifs; for example, the presence of a T residue at position 7 in the CDR3B uniquely indicated binding affinity to the epitope ELAGIGILTV (Figure 4B, rightmost column and bottom row), while the presence of an E residue at the same position indicated binding affinity to GLCTLVAML (Figure 4B, second column from the right and bottom row). To combine the individual MCMC runs containing redundant sequence information, we clustered the MCMC chains using the following method: for every pair of CDR3B sequence length and epitope sequence, we clustered the 200 MCMC chains into *Q* number of clusters, with *Q* ranging from 2 to 40; the clusters were obtained by associating each chain with its empirical amino acid distribution across MCMC samples and performing hierarchical clustering with complete linkage, defining the distance between MCMC chains as the Jenson-Shannon divergence between their corresponding empirical amino acid distributions (Methods: Model interpretation). We calculated the Silhouette score of the clustering for every value of *Q* within the specified range and chose the value of *Q* displaying the highest “peak” in Silhouette score as the final value used for clustering (Supplementary Figure S6).

Individual MCMC runs contained within the same cluster were combined, forming a single MCMC run termed the “cluster MCMC run.” We then ranked each cluster MCMC run according to a uniqueness score, such that the most unique cluster MCMC run had the highest rank (Methods: Model interpretation). The top 3 most unique cluster MCMC runs associated with CDR3B sequences of length 15 binding the top 4 epitopes are shown in Figure 4B. The top 3 most unique cluster MCMC runs for every combination of CDR3B sequence length and epitope sequence are shown in Supplementary Figure S7. Our model interpretation method identified subtle individual motifs associated with particular epitopes. For example, the CDR3B sequences of length 15 binding the epitope GILGFVFTL preferred the motifs TSS and STV at positions 7-9 (Figure 4B, first column from the left, top 2 rows), whereas CDR3B sequences of length 15 binding the epitope NLVPMVATV preferred the motifs TAA and STG at positions 7-9 (Figure 4B, second column from the left, top 2 rows). Additionally, our interpretation method identified key positions and amino acids contributing to binding. For example, having an L, A, or V residue at position 8 indicated preferential binding to ELAGIGILTV (Figure 4B, first column from the right, second row), whereas having the same residues at position 7 indicated preferential binding to NLVPMVATV (Figure 4B, second column from the left, third row). Likewise, CDR3B sequences with an S, G, or P residue at position 10 indicated preferential affinity to GLCTLVAML (Figure 4C, second column from the right, second row), and sequences displaying G, E, T, and R residues at position 6 showed preferential affinity to GILGFVFTL. These results demonstrated that our model interpretation method highlighted a wide variety of motifs that differentiated CDR3B sequences by epitope specificity. However, the diversity and large number of motifs identified by the model suggested that simple rules may not be sufficient to classify CDR3B sequences by their binding epitope.

To further interpret the learned model features, we investigated the physical and biochemical properties of preferred amino acids in CDR3B sequences by representing the empirical amino acid distributions of MCMC runs as matrices constructed by utilizing the 6-dimensional Atchley representation of amino acids (Methods: CDR3B and epitope sequence representation). Given an MCMC run associated with a CDR3B sequence of length *L* and epitope sequence, a 5 × *L* matrix was constructed by performing a weighted average of the first 5 entries of the Atchley representation of sampled amino acids at each position, with weights given by the empirical amino acid frequency from the MCMC run. Each of the 5 entries corresponded to a factor score capturing a set of amino acid chemical properties (36). Roughly, Factor 1 correlates with polarity and hydrophilicity, Factor 2 correlates with propensity for amino acids to be included in various secondary structural configurations, Factor 3 correlates with molecular size, Factor 4 anticorrelates with refractivity and heat capacity while correlating with propensity for amino acids to be included in various proteins, and Factor 5 correlates with positive charge. An entry in the matrix therefore reflected a particular set of amino acid biochemical properties at the corresponding position in the CDR3B sequence, and its extreme value signified that the embodied amino acid properties at the position influenced the predicted binding affinity. Matrices were constructed from individual MCMC runs, yielding “individual matrices,” and representative MCMC runs (Methods: Model interpretation), yielding “representative matrices.” Differences between the populations of CDR3B sequences of TCRs binding different epitopes were evident in the heatmap visualization of the representative matrices excluding the 4 conserved positions at the N-terminus and the C-terminus (Figure 4C, Supplementary Figure S8). For example, the heatmap associated with the epitope GILGFVFTL was characterized by low values for Factors 3 and 5 at position 7, suggesting that binding this epitope required small negatively charged amino acids at position 7; CDR3B sequences binding NLVPMVATV were characterized by nonpolar amino acids at positions 6-8 and amino acids with low refractivity and heat capacity at positions 6 and 8; CDR3B sequences binding GLCTLVAML were characterized by a small negatively charged amino acid at position 5 and an amino acid with high Factor 2 score at position 8; finally, CDR3B sequences binding the epitope ELAGIGILTV had similar biochemical profiles as CDR3B sequences binding the epitope NLVPMVATV, but preferred amino acids with lower Factor 2 score at position 7.

To gain a more fine-grained understanding of the chemical properties contributing to binding, we clustered the 200 individual matrices associated with each combination of CDR3B sequence length and epitope sequence. Clustering was performed using agglomerative clustering with complete linkage and with the distance metric between individual matrices defined by the Frobenius distance. As in the clustering of weblogos, we determined the number *R* of clusters by clustering the matrices with values of *R* ranging from 2 to 40, calculating the Silhouette score for each value of *R*, and choosing the value of *R* displaying the highest “peak” (Supplementary Figure S9). A single matrix, termed the “cluster matrix,” was constructed for each cluster by averaging the individual matrices contained in the cluster. The cluster matrices were then ranked according to a uniqueness score given by its average Frobenius distance to the representative matrices with the same CDR3B sequence length and alternative epitope sequence (top 3 clusters shown in Figure 4D and Supplementary Figure S10). These fine-grained heatmaps displayed unique biochemical patterns not captured by the representative heatmaps (Figure 4C). For example, while a consensus propensity for hydrophobic amino acids at positions 6-8 was evident for NLVPMVATV (Figure 4C), the refined heatmaps for NLVPMVATV revealed a preference for hydrophobic amino acids at either position 7 or 8, suggesting that only a single hydrophobic amino acid was necessary at one of these locations (Figure 4D, second column from the left). Additionally, while hydrophobic amino acids seemed to be prevalent at positions 6-8 for the vast majority of CDR3B sequences binding the top 4 epitopes, certain CDR3B sequences binding GLCTLVAML and ELAGIGILTV did not possess this property (Figure 4D). These results demonstrated that our model extracted epitope specificity through learning interpretable patterns of amino acid chemical properties in the CDR3B sequence.

### Salient positions in CDR3B sequence do not necessarily reflect physical proximity to epitope

We next sought to analyze the model interpretation results in the context of available TCR-pMHC crystal structure data. Specifically, we investigated the assertion by Glanville et al. (14) that the central positions of CDR3B sequence contacting the epitope play an important role in determining TCR-epitope specificity. To investigate this assertion, we utilized the published crystal structure data of TCR-pMHC complexes (38) restricted to HLA A*02:01 and containing one of the top 4 epitopes with the most reported interactions from the fixed dataset (GILGFVFTL, NLVPMVATV, GLCTLVAML, ELAGIGILTV). Computing the minimum distance between a given position in the CDR3B sequence and the bound epitope showed that the center of the CDR3B sequence tended to be closest to the epitope (Figure 5A and 5B; Supplementary Figure S11 and S12; Methods: TCR-pMHC structure analysis). We investigated whether the salience of a particular position inferred by our interpretation method correlated with the physical proximity of the amino acid to the epitope. We first quantified the salience of a position in the CDR3B region to be the KL divergence of the amino acid distribution derived from MCMC samples with respect to the uniform amino acid distribution (Figure 5C; Supplementary Figure S13). As the first and last four positions showed high conservation of the CASS and EQYF motifs at the N-terminus and C-terminus, respectively, across the top 4 epitopes (Figure 4A), suggesting that these residues do not play a major role in determining epitope specificity, we excluded these positions from the analysis.

**Figure 5:**
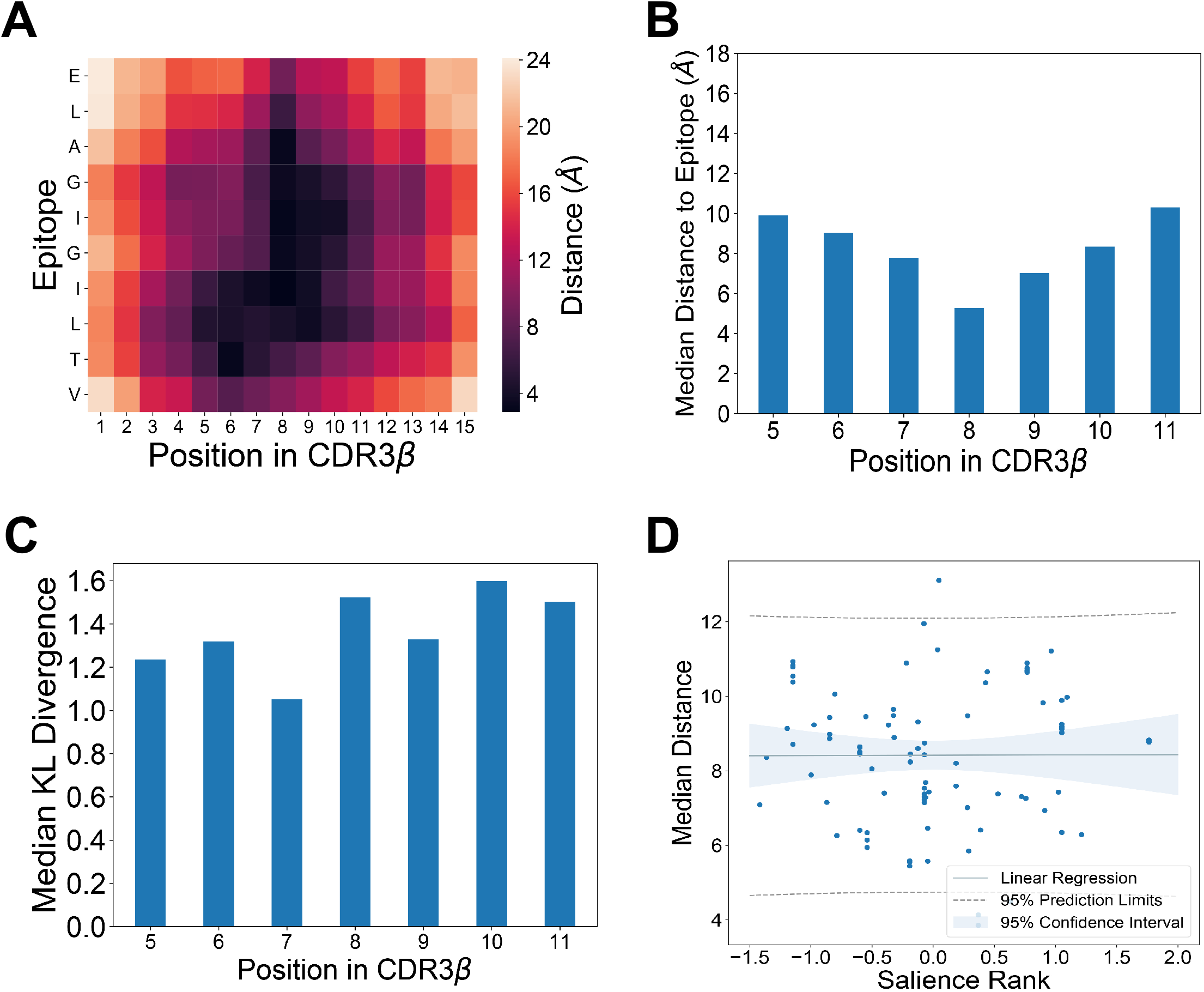
Analysis of TCR-pMHC crystal structure data reveals limited relationship between physical proximity and salience in binding specificity. **(A)** Heatmap visualizing the pairwise distance between amino acids in the CDR3B chain and the epitope ELAGIGILTV, calculated from TCR-pMHC crystal structure data (38). **(B)** Median distance to the epitope ELAGIGILTV for each position within the CDR3B sequence of length 15 (38). The middle positions of the CDR3B sequence tended to be more proximal to the epitope, with positions 8 and 9 displaying the highest proximity. **(C)** Bar chart displaying the median KL divergence of the empirical amino acid distribution from the uniform distribution at each position in the CDR3B. Median was taken across all MCMC runs associated with CDR3B sequences of length 15 binding the epitope ELAGIGILTV. **(D)** Scatterplot with linear regression fit showing statically insignificant correlation (*p* = 0.976) between the salience rank of a position according to the MCMC interpretation method and its median distance to the epitope derived from TCR-pMHC crystal structure data. Dotted lines indicate 95% prediction limit, and shaded area indicates 95% confidence interval.

We investigated the connection between the model-learned importance of a position in the CDR3B region and the physical proximity of the amino acid to the epitope according to structural data by assessing the correlation between a summary measure of position salience obtained from the model interpretation method, called the “salience rank,” and the median distance of the position to the epitope (Methods: TCR-pMHC structure analysis). Intuitively, a low salience rank for a position signified that the model interpretation method found the position to be important for epitope specificity. We did not detect a statistically significant correlation (Methods: TCR-pMHC structure analysis) between the salience rank of a position and its transformed median distance to epitope (Figure 5D; Pearson correlation = 0.0032; *p* = 0.976). These results suggested that the majority of the inner amino acids of the CDR3B sequence, not just those most proximal to the bound epitope, contributed to determining the epitope specificity of TCR. Although the most proximal positions might be still important for interacting with epitopes, as evidenced by the high degree of sequence conservation at the central locations (Figure 4A), these amino acids were also conserved across CDR3B sequences of TCRs binding different epitopes, suggesting that they might mediate interactions with generic epitopes, rather than determining epitope specificity. These results indicate that TCR-epitope specificity may rely on CDR3B sequence features not necessarily most proximal to the epitope, contrary to the common assumption.

## DISCUSSION

We have introduced a hybrid convolutional neural network model to represent TCR and epitope sequences in a shared latent space and study TCR-epitope binding specificity. Inspired by ideas from multimodal machine learning (28) and metric learning (25,26), our method can be viewed as modeling the TCR-epitope binding interactions using a Gaussian mixture model in the latent space. Evaluating the performance of this model on multiple subsets of a dataset consisting of positive binding pairs of CDR3B and epitope sequences aggregated from four popular databases (29–31,34) and nonbinding CDR3B sequences from a large cohort of healthy volunteers (35), we have shown that our model achieves similar performance, and in some cases superior performance, compared to other state-of-the-art machine learning methods for studying TCR-epitope binding (20,23). Additionally, our hybrid neural network model provides an interpretable latent space in which CDR3B sequences are embedded as points and epitope sequences are mapped to Gaussian distributions, giving an intuitive spatial representation of the decision boundaries drawn by the model. Furthermore, we have identified key differences in preprocessing of CDR3B sequences in public databases and shown that ignoring such differences may inadvertently force machine learning models to learn to utilize epitope-specific preprocessing artifacts to predict TCR-epitope binding, thereby potentially inflating the assessment of model performance.

Interpreting the learned features of our neural network model through an MCMC sampling approach has enabled the identification of epitope-specific motifs and biochemical properties of binding CDR3B sequences. The results highlight that a given epitope is associated with a large number of diverse CDR3B motifs, rather than a few monolithic motifs. This diversity emphasizes the difficulty of TCR-pMHC binding prediction, suggesting that TCR-pMHC binding may depend on many complicated factors arising from interactions between the CDR3B region and the epitope. Despite these challenges, our analysis has identified several informative trends in CD3B sequences, such as the increased sensitivity of binding affinity to changes in discriminatory amino acids. Integrating these results with known crystal structures of TCR-pMHC complexes (38) has shed physical insight into the predictive features learned by the model, demonstrating that the salient positions within the CDR3B sequence may not necessarily be the closest to the binding epitope. The salience assigned by our model interpretation method to the amino acids more distant to the epitope suggests that these residues may still contribute to epitope specificity by weakly interacting with the epitope themselves or providing specific structural support to the amino acids directly contacting the epitope. Our work thus advances two key tasks associated with TCR-pMHC prediction and highlights several biochemical properties of the discriminatory amino acids learned by our model.

## Author Contributions

J.S.S. conceived and supervised the project; A.L. developed the algorithms and analyzed the data; J.R.L., T.M, and S.K. assisted with data analysis; J.S.S. and A.L. wrote the manuscript, and all other authors have read the manuscript and contributed to the writing.

## Data Availability

The source code is available in the GitHub repository (https://github.com/jssong-lab/TCR-Epitope-Binding)

## Supplementary Materials

The following are available online: Supplementary Methods, Supplementary Tables S1-S6, Supplementary Figures S1-S13, and References for Supplementary Materials

