## Supplementary Methods, Tables, and Figures for "Predicting TCR-epitope Binding Specificity Using Deep Metric Learning and Multimodal Learning"

**This PDF file includes:**

Supplementary Methods
Supplementary Tables S1 to S6
Supplementary Figures S1 to S13
References for Supplementary Methods, Tables and Figures

### Supplementary Methods

#### Model architecture and training

Our model consisted of two components: a CDR3B embedding network (Figure S1) that maps the Atchley representation $x$ of a CDR3B sequence to a vector $f(x)$ in a 32-dimensional latent vector space and an epitope mapping network (Figure S2) that maps the Atchley representation $y$ of an epitope sequence to the parameters $\mu(y)$ and $\sigma(y)$ specifying a Gaussian distribution over the same latent vector space. The CDR3B embedding network passes $x$, which takes the form of a 20x6 matrix, through two 1D convolutional layers and an output fully connected layer. The first convolution layer had 128 filters using 3x6 kernels, convolving the rows with a stride of 1. The second convolution layer had 64 filters using 4x6 kernels, convolving the rows with a stride of 1. Outputs of the convolutional layers were padded with zeros to maintain the same dimension as the inputs before undergoing batch normalization. The activation function used in the convolutional layers was the Rectified Linear Unit (ReLU). The output fully connected layer consisted of 32 neurons, resulting in a 32-dimensional vector representation $f(x)$ for the given CDR3B sequence $x$.

The epitope mapping network first passed input $y$, which takes the form of a 10x6 matrix, through a single 1D convolutional layer that had 64 filters using 5x6 kernels, convolving the rows with a stride of 1. The output of the convolutional layer was padded with zeros to maintain the same dimension as the input before being passed to two separate fully connected layers, each of dimension 32. The outputs of the two fully connected layers, denoted as $\mu(y)$ and $\sigma(y)$, were taken as the parameters of a multivariate Gaussian distribution with diagonal covariance matrix, with $\mu(y)$ taken to be the mean vector and $\sigma(y)$ corresponding to the diagonal entries of the diagonal covariance matrix $\Sigma(y)$. Given CDR3B sequence $x$ and epitope sequence $y$, the unnormalized binding affinity $p$ was thus calculated by the following:

$$p(x,y) \equiv \frac{exp\left(-\frac{1}{2}\left(f(x) - \mu(y)\right)^T \Sigma(y)^{-1}\left(f(x) - \mu(y)\right)\right)}{\sqrt{(2\pi)^{32}|\Sigma(y)|}}$$

where $|\Sigma(y)|$ was the determinant of $\Sigma(y)$.

Although calculating unnormalized binding affinity involved only the CDR3B embedding network and epitope mapping network, tuning the parameters of these two networks to perform TCR-epitope prediction required the training of two additional neural networks: the Triplet Network and the modal alignment network. The Triplet Network (Figure S3) (1) was a single neural network composed of 3 CDR3B embedding networks which shared parameters and utilized Triplet Loss, a loss function that encouraged clustering of CDR3B embeddings by binding epitope. The Triplet Network took “valid triplets” of CDR3B sequences as input, with valid triplets defined as a set of three CDR3B sequences in which two sequences, denoted by $x^a$ and $x^p$ and termed the “anchor” and “positive” respectively, bind the same epitope, while the third, denoted by $x^n$ and termed the “negative”, binds an alternative epitope or is derived from the negative binding set (Methods: Positive binding and negative binding datasets). Given the function $f$, which represented the embedding of CDR3B sequences to vectors in a latent space by the CDR3B embedding network, and a margin parameter $m$, the Triplet Network calculated the Triplet Loss $\mathcal{L}_{Triplet}$ for a valid triplet as follows:

$$\mathcal{L}_{Triplet}(x^a, x^p, x^n) \equiv Max(\|f(x^a) - f(x^p)\|^2 - \|f(x^a) - f(x^n)\|^2 + m, 0).$$

Intuitively, training the Triplet Network to minimize Triplet Loss tuned the parameters of the CDR3B embedding network to decrease the distance between the latent space embeddings of the anchor and positive and increase the distance between the latent space embeddings of the anchor and negative. The margin parameter $m$ roughly established the length scale of the clustering in latent space and was set to 0.1.

The modal alignment network took as input a pair of binding CDR3B and epitope sequences, denoted as $x$ and $y$ respectively, and passed them through their respective CDR3B embedding and epitope mapping networks (Figure S4). The corresponding loss function, the negative log likelihood loss $\mathcal{L}_{NLL}$, was defined as

$$\mathcal{L}_{NLL}(x, y) \equiv -\log\left(p(x, y)\right).$$

Intuitively, minimizing $\mathcal{L}_{NLL}$ tuned the parameters of both the CDR3B embedding network and the epitope mapping network to maximize the probability of observing $f(x)$ given the multivariate Gaussian distribution parametrized by $\mu(y)$ and $\sigma(y)$.

During model training, the training set was divided into minibatches of 128 CDR3B sequences. For each minibatch, a set of valid triplets were extracted from the $128^3$ possible triplets of CDR3B sequences and a separate set of CDR3B sequences was constructed by including only those CDR3B sequences in the minibatch from the positive binding set. Let $v$ index each triplet in the set of valid triplets, with the anchor, positive, and negative of a triplet denoted as $x_v^a$, $x_v^p$, and $x_v^n$, respectively. Let CDR3B sequences from the positive binding set and their binding epitope sequences be denoted as $x_u$ and $y_u$, respectively, with $u$ indexing each pair. The model was trained by minimizing the combined loss function $\mathcal{L}_{combined}$ for each minibatch of data, defined as

$$\mathcal{L}_{combined} \equiv \sum_{v} \mathcal{L}_{Triplet}\left(x_v^a, x_v^p, x_v^n\right) + \alpha \sum_{u} \mathcal{L}_{NLL}\left(x_u, y_u\right)$$

with $\alpha$ being the parameter that tunes the relative strength of the two loss functions. Parameter $\alpha$ was set to 0.0001. We trained the model using the Adam optimizer with learning rate $10^{-5}$. During each epoch of training, the average combined loss function for the validation set was plotted. Training was stopped at the epoch at which the lowest approximated combined loss function for the validation set was achieved. The models were constructed and trained using the python package Keras version 2.0.8.

**MCMC method of model interpretation**

The MCMC method of model interpretation, using the maximum entropy approach of (2), sampled the space of CDR3B sequences with a bias towards those sequences with higher predicted binding affinity to a given epitope. Sampling occurred by first successively proposing new sequences generated from a seed sequence through a single amino-acid mutagenesis process, whereby a position in the CDR3B sequence, excluding the 4 positions at the N- and C-terminus, was randomly chosen to have its amino acid randomly replaced, under uniform sampling of positions and amino acids. The proposed sequence was accepted as the next sampled sequence according to an acceptance probability $r$. Given the original sequence $x$, proposal sequence $x'$, and epitope sequence $y$, the acceptance probability was given by

$$r = \min\left(1, \frac{e^{-\beta E(x',y)}}{e^{-\beta E(x,y)}}\right),$$

where the energy function

$$E(x,y) \equiv -\log\left(p(x,y)\right)$$

was the negative log likelihood loss of the modal alignment network and $\beta$ was an inverse temperature parameter. Setting $\beta \ll 1$ flattened the energy landscape, ensuring that the proposal sequence was always accepted, whereas setting $\beta \gg 1$ increasingly limited the MCMC sampling to sequences near the minima of the energy landscape, corresponding to sequences with the highest predicted binding affinities. $\beta$ was chosen by running the MCMC method with $\beta$ taking values between 1 and 3 with 0.1 step size and choosing the value of $\beta$ producing weblogos (3) with coherent motifs. Since the MCMC sampling procedure did not change the length of the CDR3B sequence, all subsequently sampled CDR3B sequences derived from a seed CDR3B sequence shared the same sequence length as the seed. Therefore, seeds of different length were chosen to visualize the motifs of CDR3B sequences of varying length. For each of 12 combinations of CDR3B sequence length and epitope sequence – corresponding to CDR3B sequences of length 13, 14, and 15 binding to the top 4 epitopes with the most data, GILGFVFTL,

NLVPMVATV, GLCTLVAML, and ELAGIGILTV – we obtained 40 seed CDR3B sequences by first training the model for 120 epochs on a pair of training and test sets specified by taking the top 4 epitopes as the seen epitope set (Methods: Model evaluation for specific seen epitope sets) and taking the top 40 CDR3B sequences in the combined set of training and test data with highest predicted binding affinity to each corresponding epitope as seeds. Five MCMC chains, or runs, were initiated and simulated for 20,000 steps from each of the 40 seeds, resulting in a total of 200 MCMC runs for each combination of CDR3B sequence length and epitope sequence. MCMC runs obtained in this manner were termed “individual” MCMC runs.

In order to consolidate similar individual MCMC runs, we clustered the 200 individual MCMC runs using agglomerative clustering with complete linkage. Clustering with agglomerative clustering required a notion of distance between individual MCMC runs, which we defined as follows. Let $A_i(j)$ be the empirical probability of seeing the $j^{th}$ amino acid at position $i$ for MCMC run $A$ and $B_i(j)$ be the equivalent for MCMC run $B$. We defined the distance $D$ between runs $A$ and $B$ to be

$$D(A,B) = \sum_i JSD(A_i(\cdot)||B_i(\cdot))$$

where $JSD(A_i(\cdot)||B_i(\cdot))$ denoted the Jensen-Shannon divergence between the amino acid distributions at position $i$ for $A$ and $B$. The number $Q$ of clusters was chosen by calculating the silhouette score for $Q$ ranging in value from 2 to 40 and choosing the value $Q$ displaying the highest peak in silhouette score. MCMC runs that were contained within the same cluster were appended together to form “cluster” MCMC runs.

To rank the cluster MCMC runs according to their uniqueness, we constructed a uniqueness score $U$ for cluster MCMC runs as follows. Let $C_{kl}$ be a cluster MCMC run where $k$ indexes the epitope sequence and $l$ indexes the CDR3B sequence length. Let $E_{mn}$ be a “representative” MCMC run, constructed by appending all 200 individual MCMC runs associated with the epitope sequence indexed by $m$ and CDR3B sequence of length $n$. The uniqueness score $U(C_{kl})$ was calculated as follows:

$$U(C_{kl}) = \frac{\sum_{m \neq k, n=l} D(C_{kl}, E_{mn})}{\sum_{m \neq k, n=l} 1},$$

where $D(C_{kl}, E_{mn})$ is the distance between MCMC runs $C_{kl}$ and $E_{mn}$.

In order to measure the importance of a position in the CDR3B region for a given MCMC run, we constructed a salience metric $s$ defined as follows. First, given MCMC run $W$, let $W_i(j)$ be the empirical probability of seeing the $j^{th}$ amino acid at position $i$ for MCMC run $W$. Then, we defined

$$s(W, i) \equiv KL(W_i(\cdot) || Z(\cdot)) ,$$

where $Z(j)$ denoted the uniform distribution over amino acids $j$.

To investigate the results of the MCMC interpretation method in terms of the physical and biochemical properties of amino acids, we constructed a 5 x $L$ matrix from each individual MCMC run, termed the "individual" matrix, with the 5 rows corresponding to the 5 entries of the amino acid vector representation and the $L$ columns corresponding to the positions of the CDR3B sequence of length $L$. The individual matrix was constructed as follows. Let $F_i(j)$ be the empirical probability of seeing the $j^{th}$ amino acid at position $i$ for MCMC run $F$ and let $a_m(j)$ be the $m^{th}$ entry out of the first 5 entries of the Atchley representation of amino acid $j$. The individual matrix $G_{mi}(F)$ corresponding to MCMC run $F$ was constructed as

$$G_{mi}(F) = \sum_j F_i(j) a_m(j) .$$

To consolidate individual matrices with redundant information, we clustered the individual matrices using agglomerative clustering with the distance between individual matrices defined by their Frobenius distance. The number $R$ of clusters was chosen by calculating the silhouette score for $R$ ranging in value from 2 to 40 and choosing the value $R$ displaying the highest peak in silhouette score. Individual matrices that were contained within the same cluster were averaged together to form “cluster” matrices. To rank the cluster matrices according to their uniqueness, we constructed a uniqueness score $V$ for cluster matrices as follows. Let $H_{kl}$ be a cluster matrix where $k$ indexes the epitope sequence and $l$ indexes the CDR3B sequence length. Let $J_{mn}$ be a “representative” matrix, formed by constructing the corresponding matrix for the representative MCMC run, with epitope sequence indexed by $m$ and CDR3B sequence of length $n$. The uniqueness score $V(C_{kl})$ was calculated as follows:

$$V(H_{kl}) = \frac{\sum_{m\neq k,n=l} \|H_{kl} - J_{mn}\|_F}{\sum_{m\neq k,n=l} 1}$$

With $\| \quad \|_F$ denoting the Frobenius norm.

#### Supplementary Tables

| Amino Acid | Entry 1 | Entry 2 | Entry 3 | Entry 4 | Entry 5 | Entry 6 |
|---|---|---|---|---|---|---|
| A | 0.23693 | 0.06158 | 0.51254 | 1.00000 | 0.50432 | 1.00000 |
| C | 0.00000 | 0.55173 | 0.49612 | 0.29962 | 0.48656 | 1.00000 |
| D | 0.75394 | 0.50652 | 0.14051 | 0.50541 | 0.00000 | 1.00000 |
| E | 0.85066 | 0.01969 | 0.79381 | 0.60600 | 0.39176 | 1.00000 |
| F | 0.10618 | 0.25908 | 0.84651 | 0.46809 | 0.59521 | 1.00000 |
| G | 0.30214 | 0.88100 | 0.77511 | 0.85803 | 0.86431 | 1.00000 |
| H | 0.52899 | 0.30707 | 0.39290 | 0.17685 | 0.51539 | 1.00000 |
| I | 0.03277 | 0.27101 | 0.87705 | 0.68172 | 0.66102 | 1.00000 |
| K | 1.00000 | 0.26713 | 0.67367 | 0.50054 | 0.79655 | 1.00000 |
| L | 0.10208 | 0.14896 | 0.41428 | 0.91779 | 0.37954 | 1.00000 |
| M | 0.21424 | 0.00000 | 0.88825 | 0.30368 | 0.72553 | 1.00000 |
| N | 0.72086 | 0.65243 | 0.77116 | 0.52975 | 0.68008 | 1.00000 |
| P | 0.48267 | 1.00000 | 0.39863 | 0.68929 | 0.30135 | 1.00000 |
| Q | 0.71645 | 0.37309 | 0.22337 | 0.43943 | 0.22626 | 1.00000 |
| R | 0.90769 | 0.40749 | 0.79700 | 0.69443 | 1.00000 | 1.00000 |
| S | 0.35129 | 0.81082 | 0.00000 | 0.75663 | 0.09692 | 1.00000 |
| T | 0.41304 | 0.51318 | 0.88749 | 0.82098 | 0.74198 | 1.00000 |
| V | 0.00189 | 0.34535 | 0.53659 | 0.91130 | 0.32253 | 1.00000 |
| W | 0.23566 | 0.42524 | 0.69136 | 0.00000 | 0.49813 | 1.00000 |
| Y | 0.50504 | 0.65298 | 1.00000 | 0.34884 | 0.77439 | 1.00000 |
| - | 0.50000 | 0.50000 | 0.50000 | 0.50000 | 0.50000 | 0.00000 |

**Table S1: The Atchley representations of the amino acids.** Entries 1-5 correspond to the min-max scaled factor scores obtained from (4), while entry 6 corresponds to an indicator variable denoting presence of a real amino acid. The amino acid symbol “-“ represents the placeholder amino acid used to pad sequences to a predefined length.

| Epitope | Number of TCRs |
|---|---|
| GILGFVFTL | 4200 |
| NLVPMVATV | 3761 |
| GLCTLVAML | 999 |
| ELAGIGILTV | 764 |
| LLWNGPMAV | 423 |
| EAAGIGILTV | 272 |
| CINGVCWTV | 112 |
| AMFWSVPTV | 110 |
| FLYNLLTRV | 96 |
| YVLDHLIVV | 79 |
| NLNCCSVPV | 67 |
| LLFGYPVYV | 62 |
| VLFGLGFAI | 61 |
| KMVAVFYTT | 50 |
| VVLSWAPPV | 48 |
| RTLNAWVKV | 46 |
| VVMSWAPPV | 44 |
| ILTGLNYEV | 42 |
| SLFNTVATLY | 38 |
| SLYNTVATL | 35 |
| KLMNIQQKL | 35 |
| FLASKIGRLV | 34 |
| ILTGLNYEA | 33 |
| KLSALGINAV | 32 |
| FLYALALLL | 28 |

**Table S2: The top 25 epitopes with the most recorded instances of CDR3B binding interactions in the fixed dataset**. The entries are ranked in descending order.

| N Terminus | | | | C Terminus | | | |
|---|---|---|---|---|---|---|---|
| Proper TCRs | | Improper TCRs | | Proper TCRs | | Improper TCRs | |
| Sequence | Count | Sequence | Count | Sequence | Count | Sequence | Count |
| ASS | 5241 | ASS | 5945 | EQY | 1363 | EQY | 1709 |
| SAR | 260 | SAR | 360 | NEQ | 1149 | TQY | 1498 |
| ASR | 250 | ASR | 331 | TQY | 986 | GYT | 922 |
| AST | 193 | ATS | 227 | EQF | 894 | PQH | 621 |
| ATS | 170 | AST | 163 | TEA | 715 | PLH | 187 |
| AWS | 103 | AWS | 117 | GYT | 672 | TIY | 168 |
| ASG | 103 | ASG | 112 | GEL | 632 | IQY | 109 |
| SVE | 85 | SVE | 95 | EAF | 632 | VLT | 105 |
| AIS | 58 | ASN | 83 | PQH | 498 | EAF | 92 |
| SAS | 55 | AIS | 71 | ELF | 432 | ELF | 84 |
| SVG | 49 | SAS | 60 | EKL | 226 | EQF | 79 |
| ASN | 46 | ASA | 42 | TIY | 166 | GEL | 60 |
| ASA | 35 | ASK | 41 | KLF | 154 | DQY | 39 |
| SAP | 30 | ASI | 39 | PLH | 143 | NEQ | 35 |
| SAT | 24 | SVG | 36 | DEQ | 123 | TEA | 33 |
| ASK | 24 | SVD | 33 | GEQ | 78 | RLT | 22 |
| SVP | 24 | SAP | 32 | VLT | 63 | YGY | 22 |
| ASI | 21 | SAT | 30 | IQY | 62 | SYT | 19 |
| ASM | 20 | ASL | 29 | AEA | 59 | EKL | 18 |
| SVD | 18 | ANS | 25 | YEQ | 43 | DTQ | 16 |

**Table S3: The number of occurrences of each 3-mer at the N-terminus and C-terminus for native proper and improper TCRs.** Conserved C at N-terminus and conserved F at C-terminus were excluded in the analysis.

| | Top 2 Classes | | Top 3 Classes | | Top 4 Classes | | Top 5 Classes | |
|---|---|---|---|---|---|---|---|---|
| Epitope | Mean AUC | Std AUC | Mean AUC | Std AUC | Mean AUC | Std AUC | Mean AUC | Std AUC |
| GILGFVFTL | 0.7757 | 0.0032 | 0.7639 | 0.0099 | 0.7588 | 0.0084 | 0.7538 | 0.0067 |
| NLVPMVATV | 0.7678 | 0.0053 | 0.7612 | 0.0139 | 0.7410 | 0.0094 | 0.7261 | 0.0168 |
| GLCTLVAML | | | 0.6616 | 0.0244 | 0.6818 | 0.0202 | 0.6801 | 0.0338 |
| ELAGIGILTV | | | | | 0.7558 | 0.0147 | 0.7802 | 0.0310 |
| LLWNGPMAV | | | | | | | 0.8010 | 0.0143 |

**Table S4: Table of the mean and standard deviation of Area Under the ROC Curve (AUC) scores of the model on Task 2 for each epitope class using the fixed dataset.** Performance was assessed using 5-fold cross validation. The seen epitope set ranged from the top 2 epitopes to the top 5 epitopes with the largest number of reported interactions (Table S2).

| | Top 5 Classes | | Top 10 Classes | | Top 15 Classes | | Top 20 Classes | |
|---|---|---|---|---|---|---|---|---|
| Epitope | Mean AUC | Std AUC | Mean AUC | Std AUC | Mean AUC | Std AUC | Mean AUC | Std AUC |
| EAAGIGILTV | 0.7689 | 0.0275 | 0.7969 | 0.0173 | 0.7801 | 0.0206 | 0.7827 | 0.0295 |
| CINGVCWTV | 0.5828 | 0.0978 | 0.6510 | 0.0655 | 0.6459 | 0.0578 | 0.6236 | 0.0852 |
| AMFWSVPTV | 0.7990 | 0.0815 | 0.8228 | 0.0762 | 0.8062 | 0.0698 | 0.8017 | 0.1019 |
| FLYNLLTRV | 0.7936 | 0.0322 | 0.7974 | 0.0355 | 0.7669 | 0.0641 | 0.8177 | 0.0542 |
| YVLDHLIVV | 0.6231 | 0.0539 | 0.5889 | 0.0669 | 0.5841 | 0.0505 | 0.6269 | 0.1207 |
| NLNCCSVPV | | | 0.6179 | 0.0909 | 0.6947 | 0.0472 | 0.7140 | 0.1150 |
| LLFGYPVYV | | | 0.7275 | 0.0690 | 0.5770 | 0.1115 | 0.6364 | 0.0775 |
| VLFGLGFAI | | | 0.4875 | 0.0847 | 0.4394 | 0.0749 | 0.4673 | 0.0547 |
| KMVAVFYTT | | | 0.5567 | 0.0646 | 0.5011 | 0.1538 | 0.5590 | 0.0544 |
| VVLSWAPPV | | | 0.5940 | 0.0635 | 0.8649 | 0.0535 | 0.9115 | 0.0801 |
| RTLNAWVKV | | | | | 0.5861 | 0.0450 | 0.6254 | 0.1440 |
| VVMSWAPPV | | | | | 0.8895 | 0.0772 | 0.9464 | 0.0662 |
| ILTGLNYEV | | | | | 0.6601 | 0.0665 | 0.8828 | 0.0755 |
| SLFNTVATLY | | | | | 0.5814 | 0.0783 | 0.6834 | 0.0647 |
| SLYNTVATL | | | | | 0.6763 | 0.1192 | 0.7068 | 0.0766 |
| KLMNIQQKL | | | | | | | 0.8875 | 0.0912 |
| FLASKIGRLV | | | | | | | 0.5481 | 0.1252 |
| ILTGLNYEA | | | | | | | 0.7117 | 0.1224 |
| KLSALGINAV | | | | | | | 0.9515 | 0.0720 |
| FLYALALLL | | | | | | | 0.8583 | 0.0667 |

**Table S5: Table of the mean and standard deviation of Area Under the ROC Curve (AUC) scores of the model on Task 2 for each epitope class using the "reduced fixed dataset".** The reduced fixed dataset (Methods: Model evaluation for specific seen epitope sets) was constructed from the fixed dataset by removing CDR3B sequences binding the top 5 epitopes with the largest number of reported interactions (Table S2). Performance was assessed using 5-fold cross validation. Analysis included seen epitope sets corresponding to the top 5, 10, 15, and 20 epitopes with the largest number of reported interactions in the reduced fixed dataset.

| Peptide | ERGO | | TCRGP | | Our Model |
|---|---|---|---|---|---|
| | McPAS + VDJdb | | (B,3), LOSO | (B,3), unique LOO | |
| | AE | LSTM | | | |
| GILGFVFTL | 0.725 | 0.712 | 0.818 | 0.822 | 0.662 |
| NLVPMVATV | 0.624 | 0.632 | 0.587 | 0.651 | 0.761 |
| GLCTLVAML | 0.708 | 0.686 | 0.782 | 0.852 | 0.7639 |

**Table S6: Comparison of our model with the ERGO model (5) and the TCR Gaussian Process (TCRGP) model (6).** Our model and TCRGP were evaluated on Task 1, while ERGO was evaluated on a classification task similar to Task 1, which they term single peptide binding (SPB). Results shown for ERGO correspond to evaluation of the model on a combined set of McPAS (7) and VDJdb (8) data. Results shown for TCRGP are from evaluating the model only on the CDR3B data, referred to as (B,3), from (9). Our model was evaluated on a combined set of McPAS (7), VDJdb (8), IEDB (10), and PIRD (11) data. Some of the difference in the performance of TCRGP compared to ERGO and our approach may be attributed to the difference in the utilized CDR3B training and test data.

#### Supplementary Figures

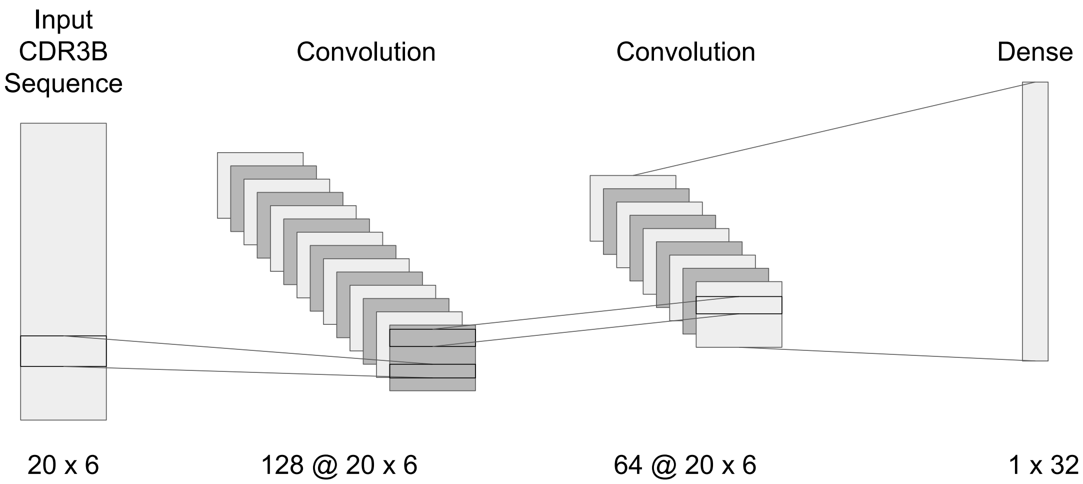


**Figure S1**: **CDR3B embedding network architecture**. A 20 x 6 Atchley representation of a CDR3B sequence is passed through two 1D convolutional layers with filter sizes {128, 64}, stride {1, 1}, and kernel sizes {3x6, 4x6}, respectively, before being passed to a dense layer of dimension 1 x 32 (Supplementary Methods: Model architecture and training). The output of the dense layer is taken to be the latent space representation of the CDR3B sequence.

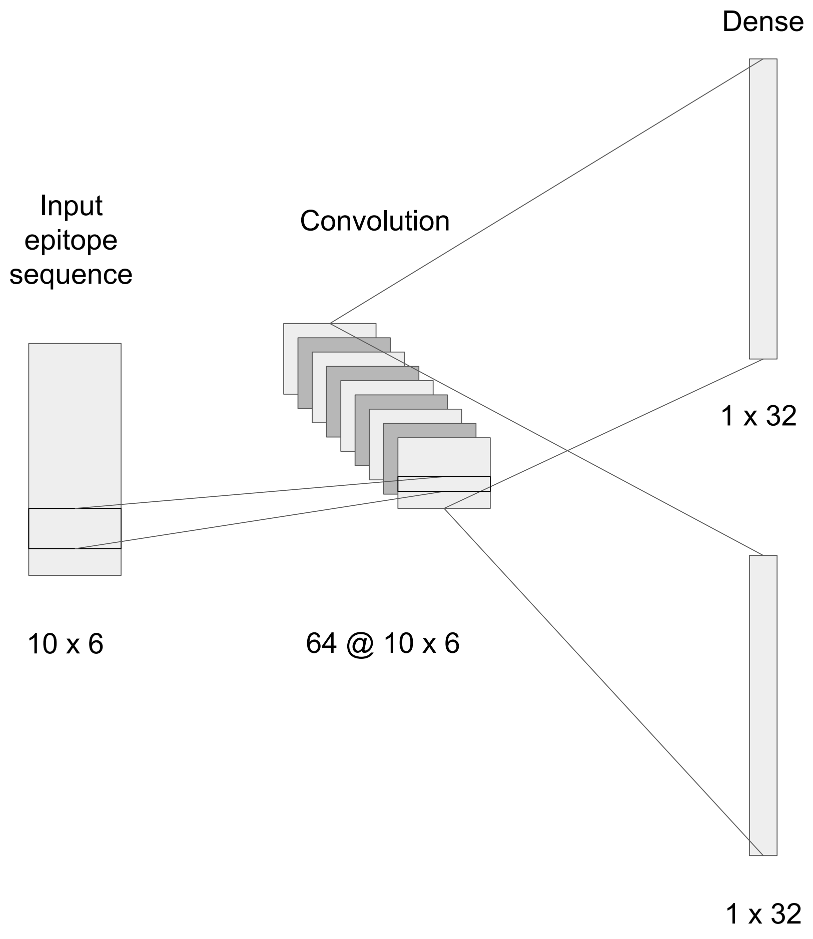


**Figure S2: Epitope mapping network architecture.** A 10 x 6 Atchley representation of an epitope sequence is passed through a single 1D convolutional layer with filter size 64, kernel size 5x6, and stride 1 before being passed to two dense layers, each of dimension 1 x 32 (Supplementary Methods: Model architecture and training). The output of the two dense layers are taken to be the parameters of a multivariate Gaussian distribution in latent space.

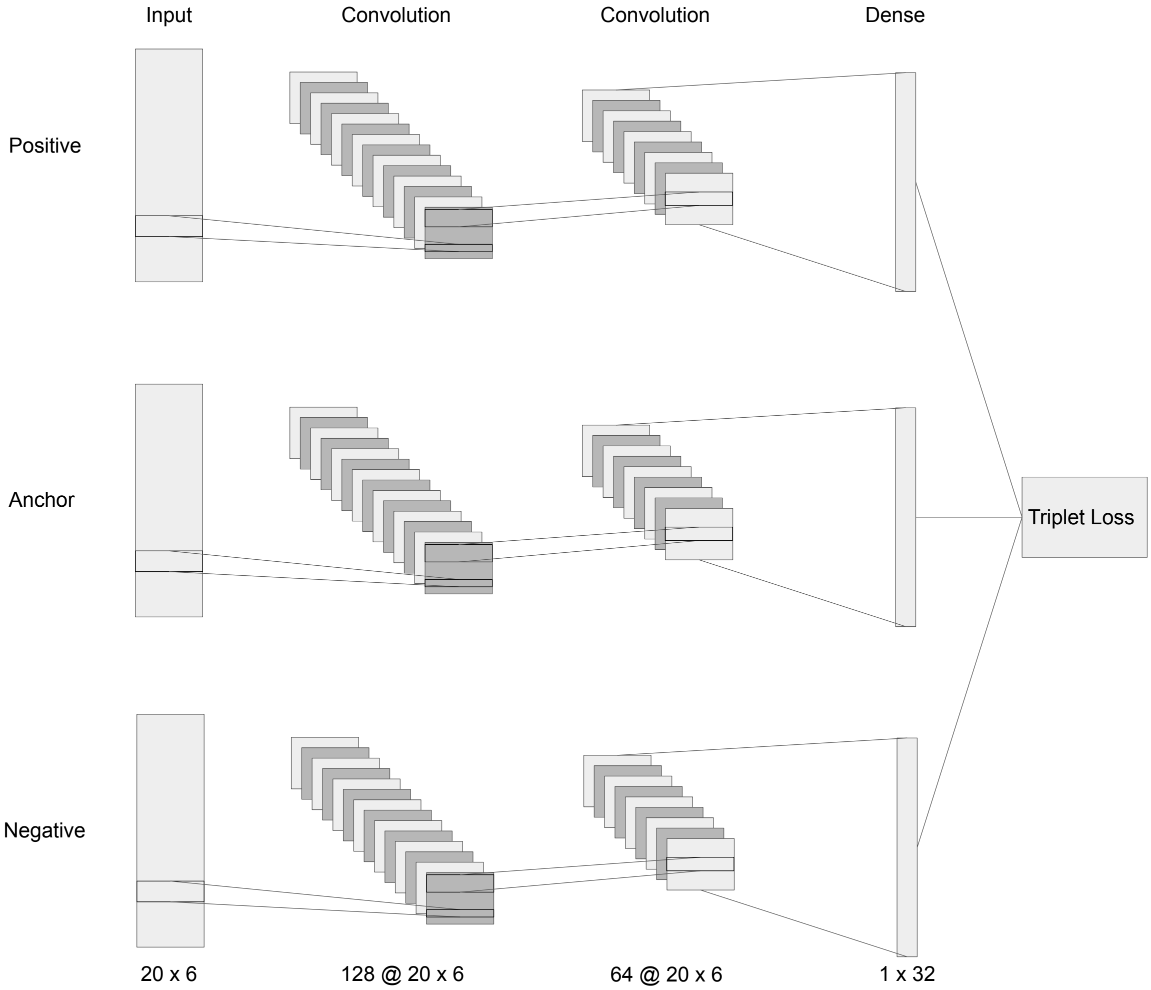


**Figure S3: Triplet Network architecture.** Three CDR3B sequences corresponding to anchor, negative, and positive sequences are each passed through the CDR3B embedding network, sharing the same parameters. The Triplet Loss is calculated from the latent space representations and used to force CDR3B sequences to cluster in latent space.

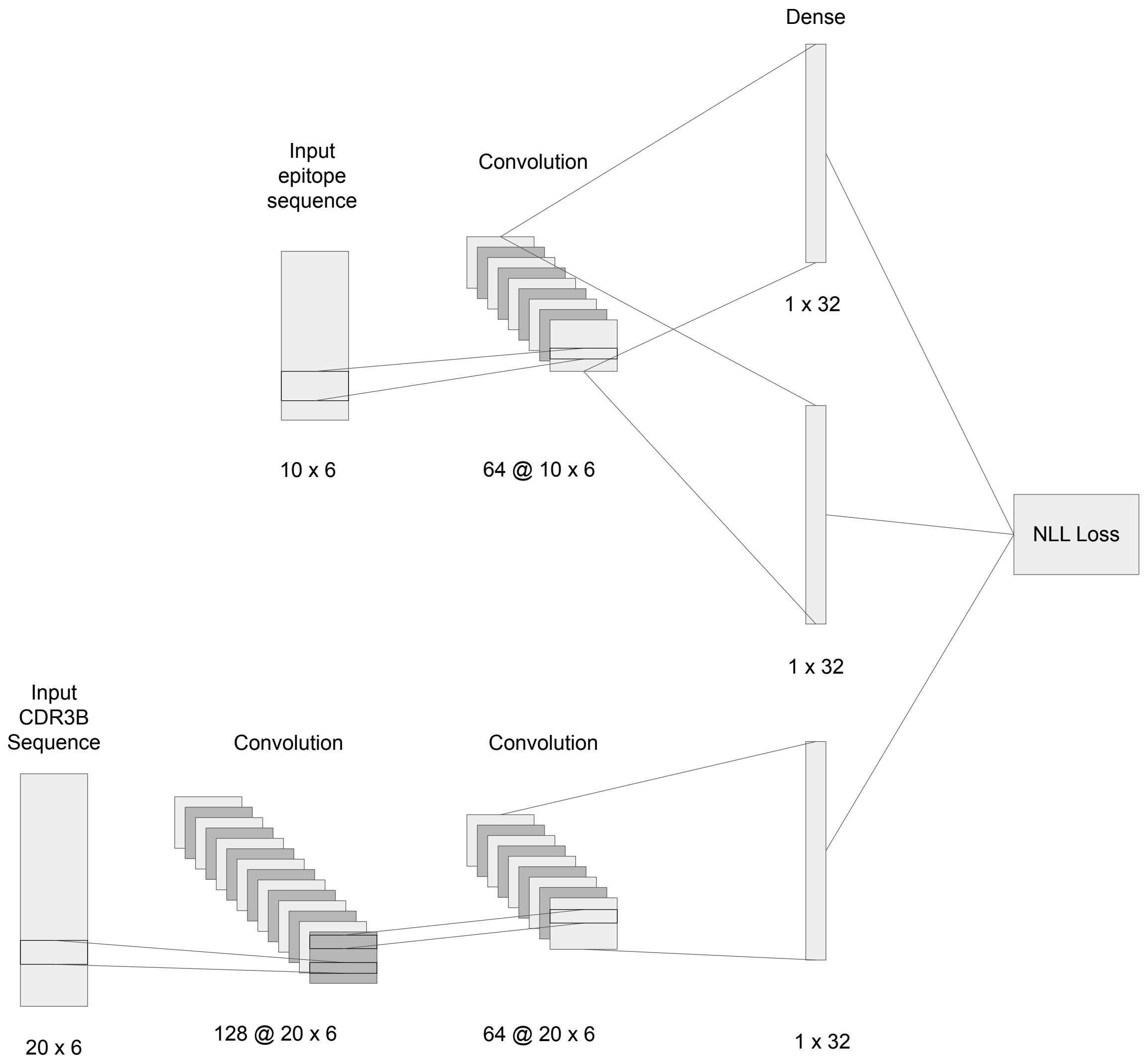


**Figure S4: Modal alignment network architecture.** The unnormalized binding affinity of a CDR3B sequence and epitope sequence is found by calculating the probability density of the point in latent space representing the CDR3B sequence given by the Gaussian distribution corresponding to the epitope sequence.

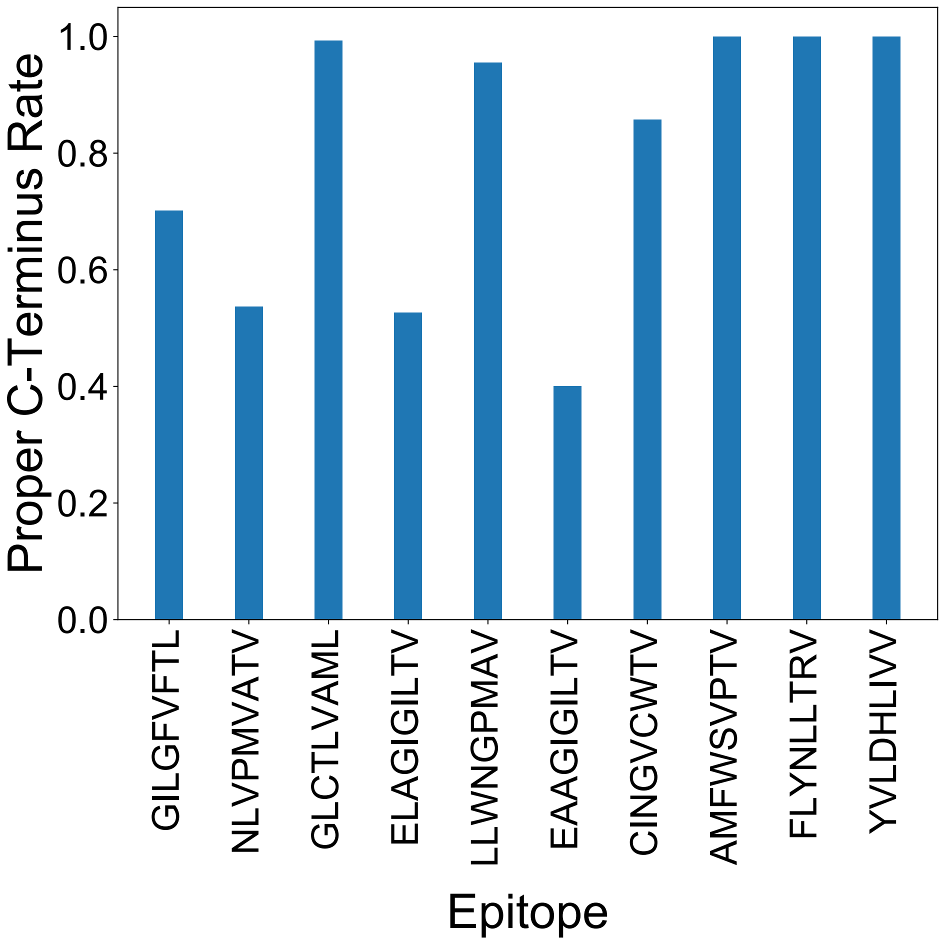


**Figure S5: Proportion of CDR3B sequences derived from the unfixed dataset displaying proper C-terminus, broken down by binding epitope.**

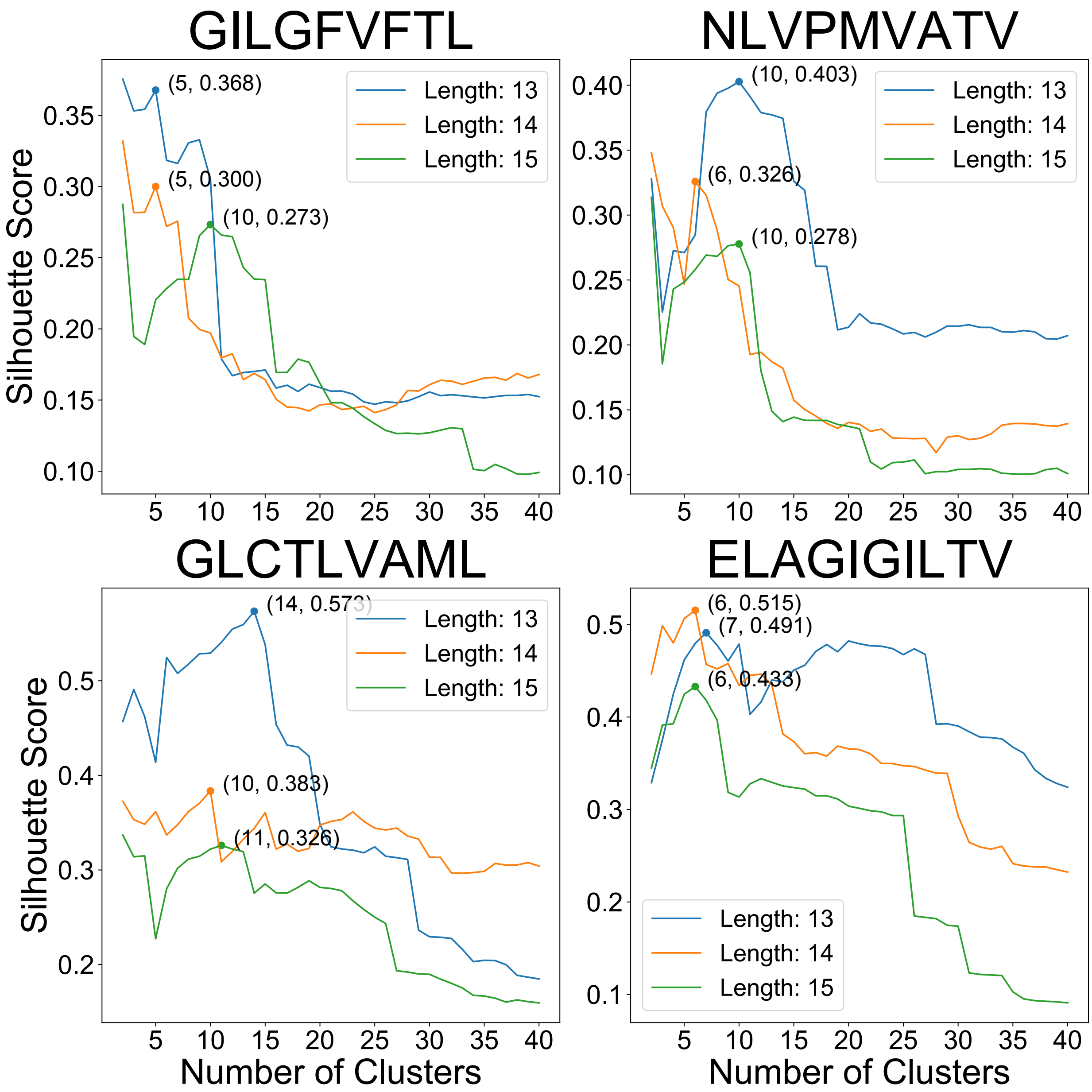


**Figure S6: Plots of Silhouette scores, obtained by clustering individual MCMC runs, with respect to number of clusters.** Final number of clusters chosen is marked by a point with coordinates.

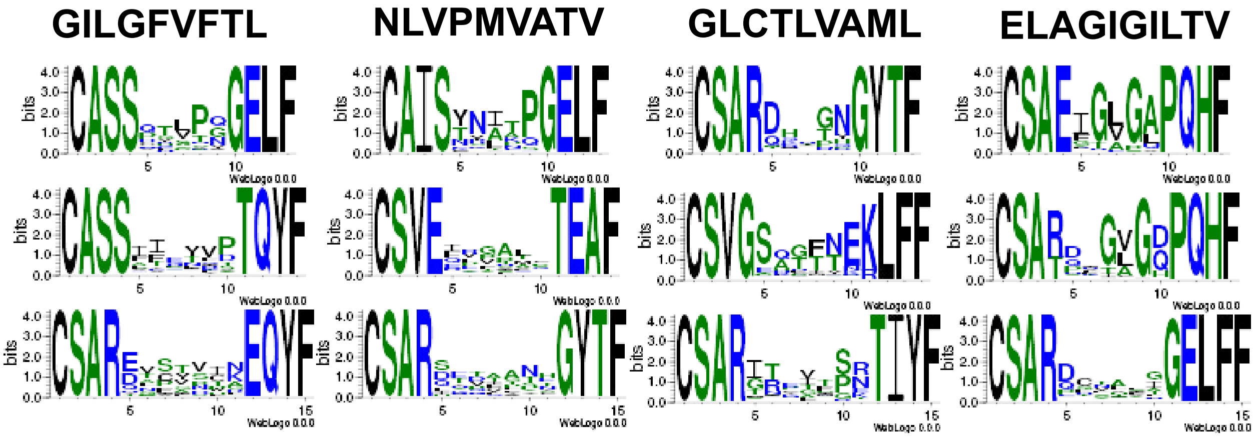


**Figure S7: Weblogos visualizing the most unique "cluster" MCMC run for each combination of CDR3B sequence length and epitope sequence.** Logo size at each position represents the extent of sequence conservation.

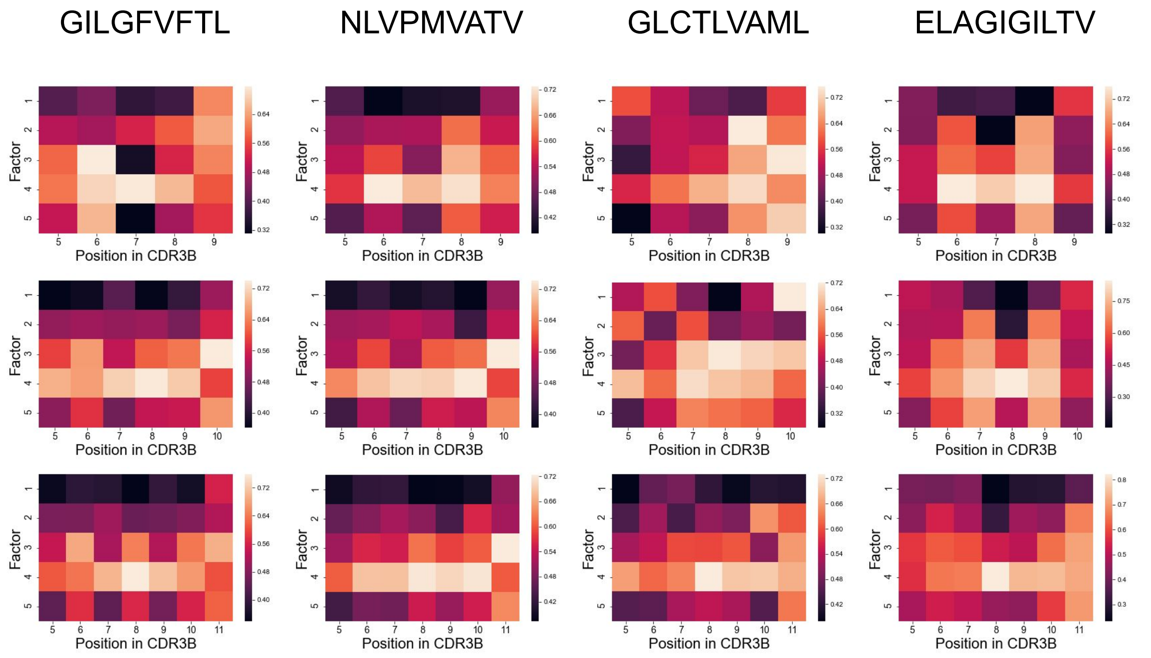


**Figure S8: Heatmaps visualizing the "representative" matrix for each combination of CDR3B sequence length and epitope sequence.** The four conserved positions at the N- and C- terminus are excluded for clarity.

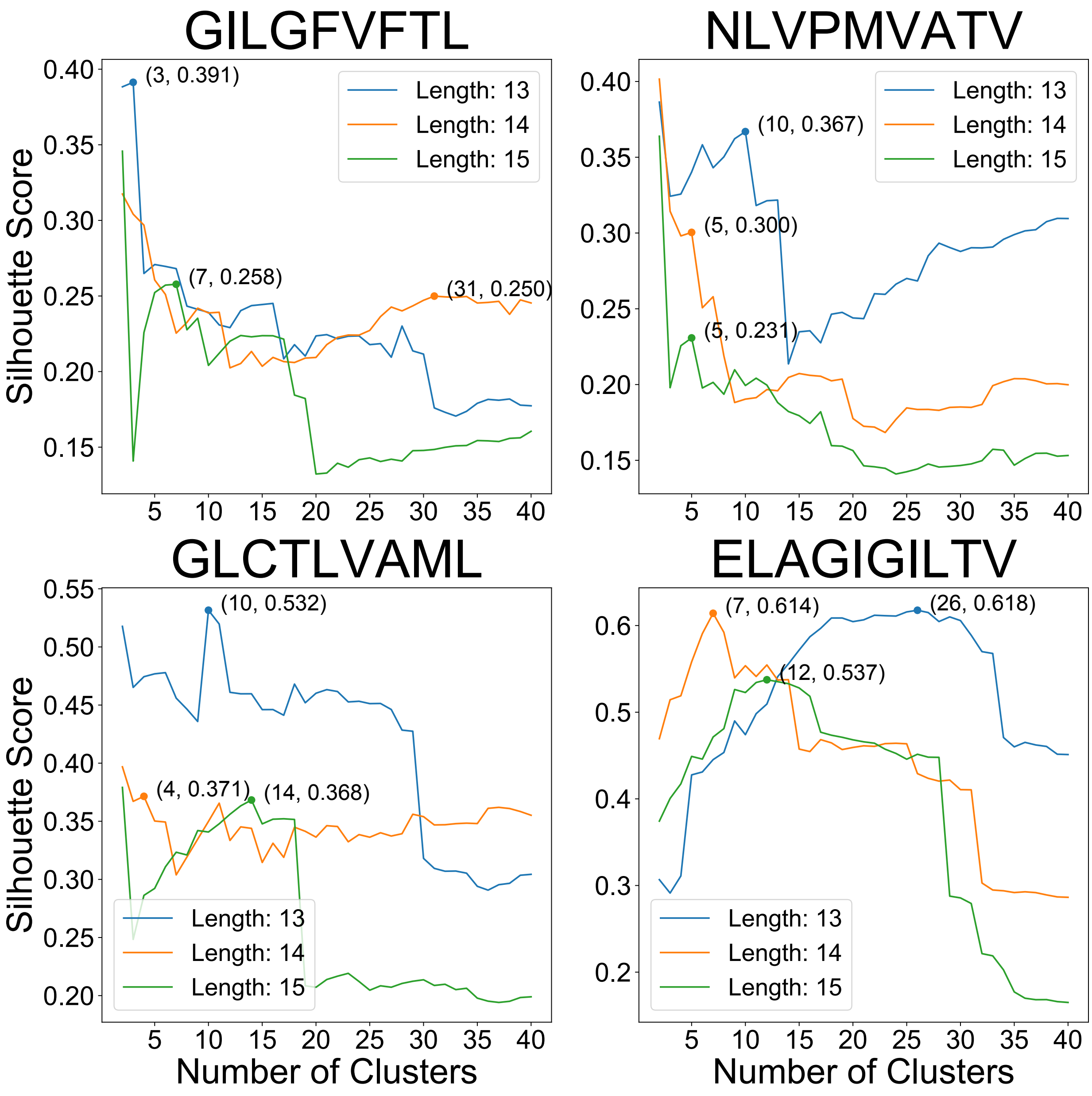


**Figure S9: Plots of Silhouette scores, obtained by clustering individual matrices, with respect to number of clusters.** Final number of clusters chosen is marked by a point with coordinates.

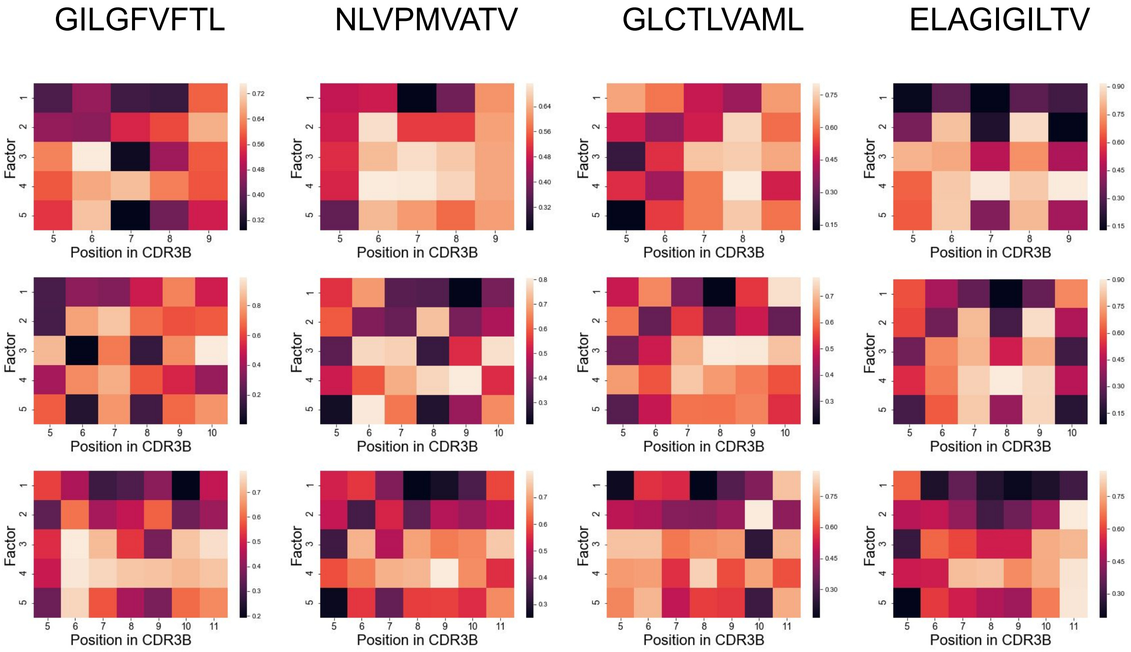


**Figure S10: Heatmaps visualizing the most unique "cluster" matrix for each combination of CDR3B sequence length and epitope sequence.** The four conserved positions at the N- and C- terminus are excluded for clarity.

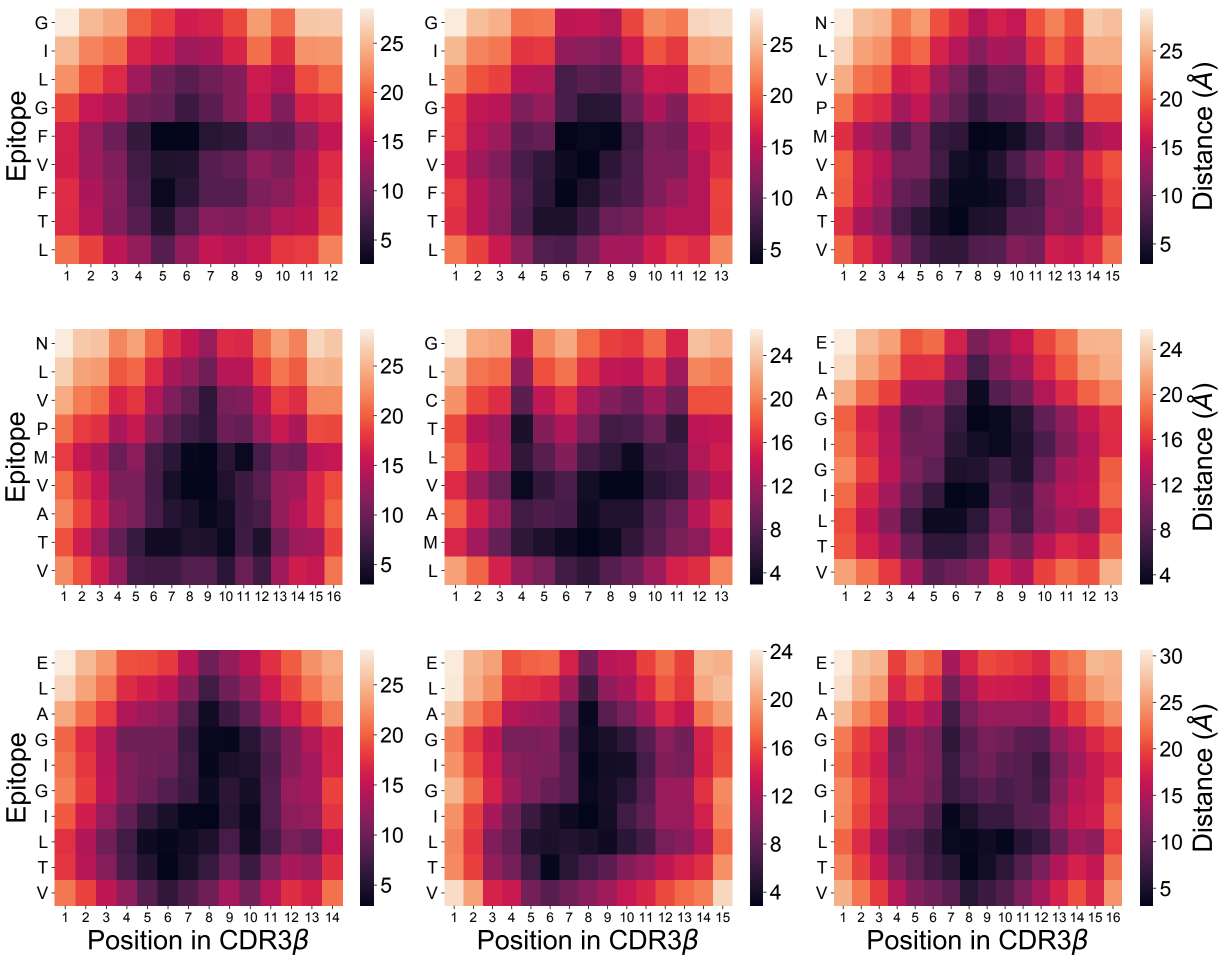


**Figure S11: Heatmaps visualizing the pairwise physical distance between amino acids in the CDR3B sequence and amino acids in the epitope sequence.** The distance was calculated from TCR-pMHC crystal structures (12) (Methods: TCR-pMHC structure analysis). Each heatmap corresponds to a specific combination of epitope and fixed CDR3B sequence length.

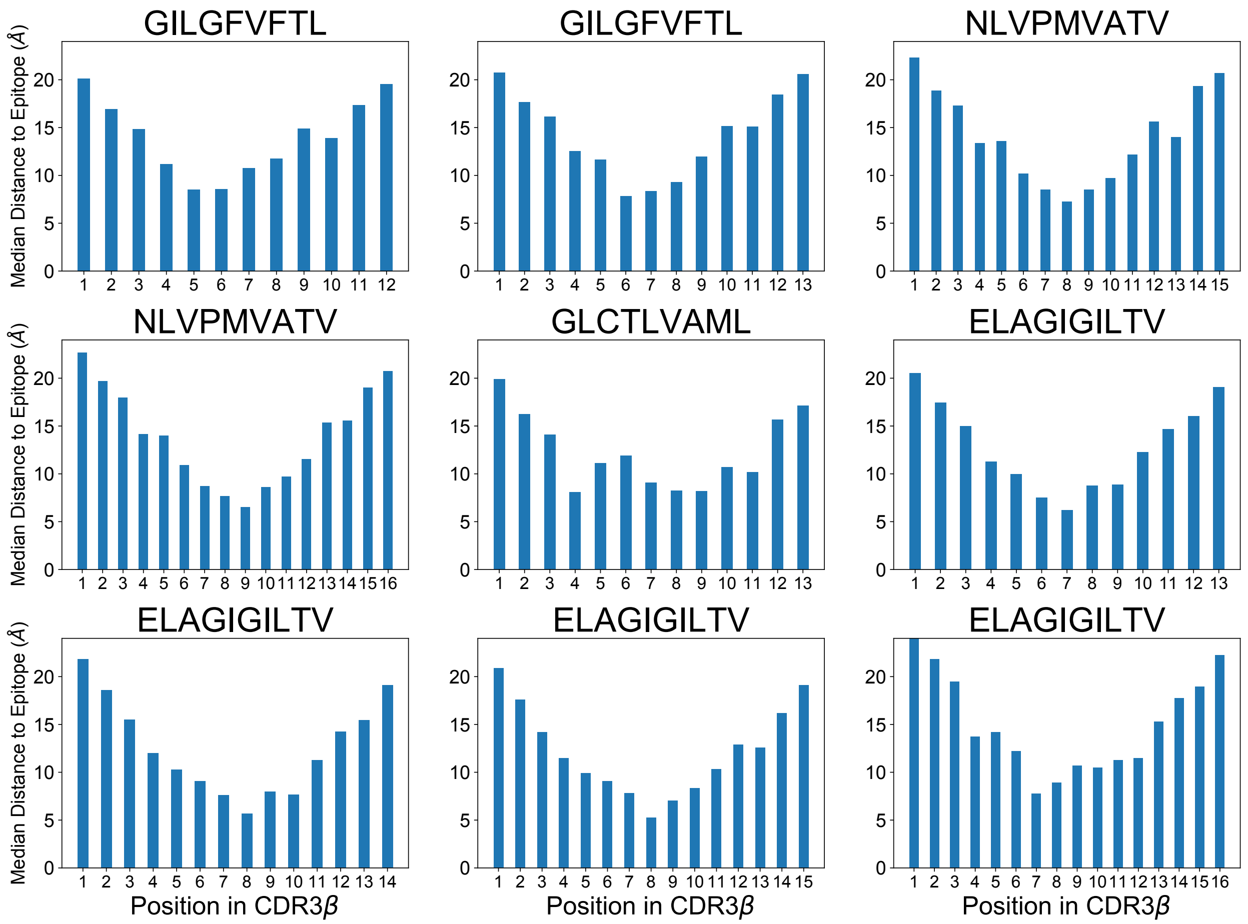


**Figure S12: Bar charts showing the median physical distance of the amino acid at each position within the CDR3B sequence to the amino acids of the epitope.** The distance was calculated from TCR-pMHC crystal structures (12) (Methods: TCR-pMHC structure analysis). Each bar chart corresponds to a specific combination of epitope and fixed CDR3B sequence length.

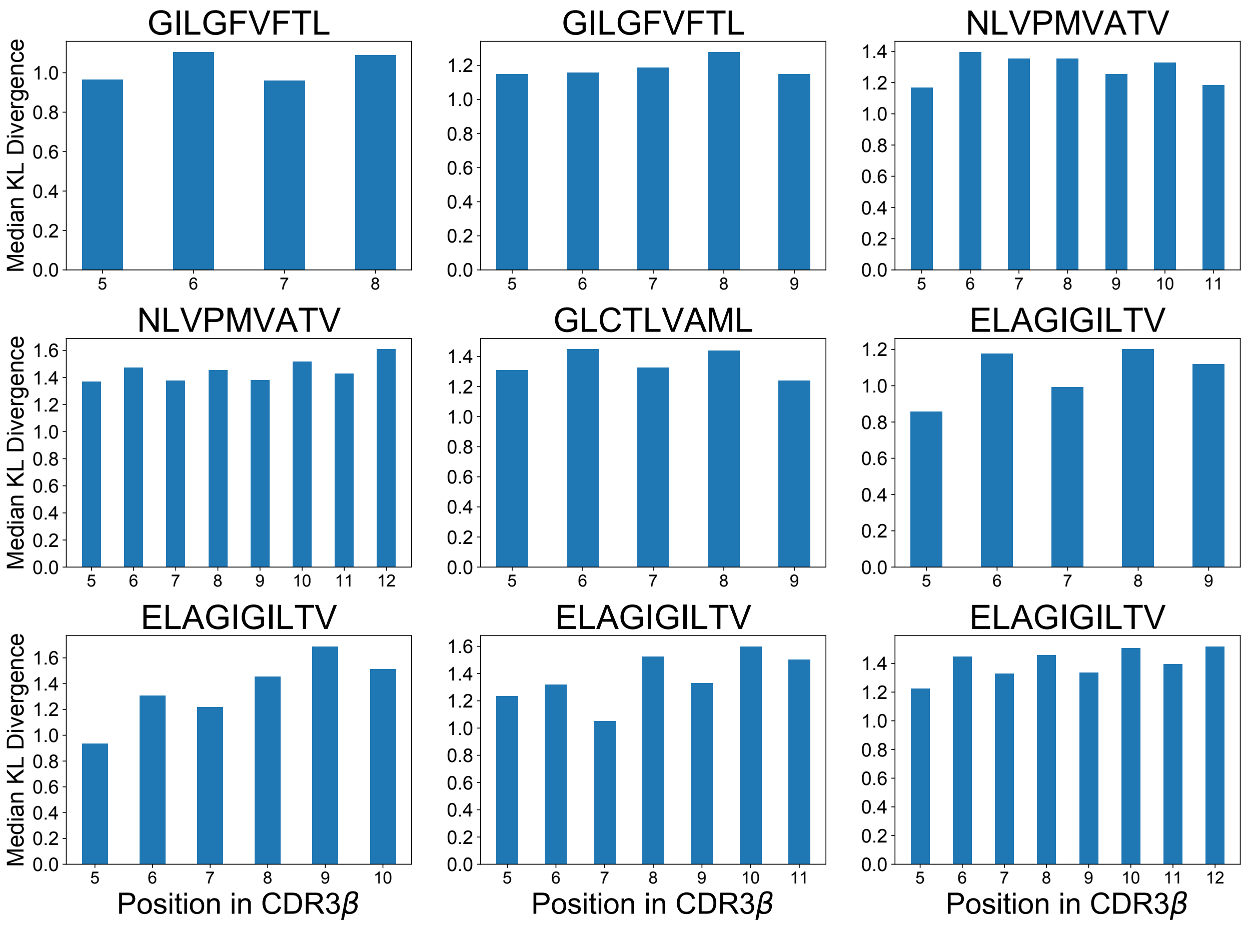


**Figure S13: Matrix of bar charts showing the median KL divergence at each position within the CDR3B sequence derived from the MCMC interpretation method.** Each bar chart corresponds to a specific combination of epitope and fixed CDR3B sequence length. The four conserved positions at the N- and C-terminus are excluded.
